# Spatiotemporal profiling reveals the role of inflammatory niche in driving prostate cancer

**DOI:** 10.64898/2026.04.19.719485

**Authors:** Abbas Nazir, Hanchen Wang, Ziyu Lu, Jeff Lau, Frank Peale, Raj Jesudason, Kelli A Connolly, Zaneta Andrusivova, Julia Lau, Sarah Gierke, Linna Peng, Sara Chan, Jian Jiang, Sandra Rost, Eric Lubeck, Marco De Simone, Bence Daniel, Lisa M. McGinnis, Danilo Maddalo, Nikhil S Joshi, Levi A Garraway, Aviv Regev

## Abstract

Prostate cancer (PCa) is a lethal malignancy that displays profound resistance to immune checkpoint blockade (ICB), via mechanisms that are poorly understood. Here, we investigate the causes of CD8 T cell exhaustion and mechanisms of tumor progression in a PCa animal model, by single cell and spatial profiling, along a time course, following orthotopic transplantation of RB1/TP53/PTEN-deficient mouse organoids, competent to express neoantigens. The resulting tumors were castration resistant, consisting of largely basal and L2 malignant cells with upregulated inflammatory gene programs, and a specific spatial distribution of macrophages, cancer associated fibroblast (CAF) subtypes, and CD8 T-cells that was not previously reported. Using Zman-seq, we demonstrate that the effector function of tumor-infiltrating CD8 T cells was rapidly impaired as early as 24hrs after their infiltration, likely driven by signals from proinflammatory macrophages, *Ccl2*-*Jak2*^+^ inflammatory CAFs, and malignant basal cells, thus driving resistance to ICB. Interestingly, dual blockade of JAK1/2 and PD1 induced potent anti-tumor effects in tumor epithelial cells, decreased malignant epithelial cells and pro-inflammatory macrophages, and increased the proportion of normal (*Pi16*^+^) fibroblasts in the TME. Our results underscore the therapeutic potential of targeting JAK1/2 to enhance the efficacy of ICB, providing a rationale for clinical investigation of this combination in PCa.

## Introduction

Prostate cancer is the second leading cause of cancer-related mortality in men (*1*). While androgen deprivation therapy and chemotherapy can produce initial responses, many patients eventually develop resistance and transition to castration resistant prostate cancer (CRPC), an aggressive, androgen-independent form of prostate cancer (*2–4*). CRPC is a heterogenous disease state, and reactivation of androgen receptor (AR) signaling is the most prevalent resistance mechanism (*2, 5*). RB1 and TP53 inactivating alterations are enriched in CRPC (*6, 7*), along with PTEN loss, a common event across prostate cancer (*6*).

Immune checkpoint blockade (ICB) therapies have revolutionized the treatment of many solid tumors (*8–10*), but their efficacy in prostate cancer has been limited. While the mechanistic basis of this resistance remains unclear, it has been broadly attributed to an unspecified immunosuppressive tumor microenvironment (TME) (*11–13*). Prostate cancer tumorigenesis involves complex interactions between mutation-bearing epithelial cells and TME cells, including macrophages, CD8 T cells, and cancer associated fibroblasts (CAFs) (*14–16*). Animal models can shed light on such interactions, but poor immunogenicity and lack of neoantigens in the tumor cells of existing preclinical prostate models has limited the study of immune cell biology in the TME. There is thus a need for novel preclinical models that recapitulate T cell biology, as seen in prostate cancer patients, and help characterize the molecular and cellular state changes within this heterogenous TME.

Here, we report a novel mouse prostate tumor model, where *Rb1^-/-^; Trp53^-/-^; Pten^-/-^* organoids, engineered from NINJA mice (*17*), are orthotopically transplanted in an immunocompetent mouse prostate, to understand tumor progression and CD8 T cell dysfunction at various stages of the disease. The NINJA mouse enables the inducible, *de novo* expression of defined neoantigens, LCMV-derived peptides GP33-43 and GP66-77, knocked-in at the *Rosa26* locus. This provides a platform to study tumor–immune interactions and the fate of tumor specific T cells. We comprehensively characterize the model by spatial transcriptomics (Xenium, 5,100 genes), to generate a high-resolution molecular map of cell and tissue organization at various stages of the disease. We further determined the kinetics of reprogramming of infiltrating immune cells by the prostate TME by time-resolved scRNA-seq using Zman-seq (*18*). The spatiotemporal patterns led us to hypothesize – and show – that PD1 blockade together with JAK1/2 inhibition leads to potent anti-tumor effects in this prostate cancer model. Our findings highlight a promising avenue for therapy, and a systematic approach that can be applied in other tumor models.

## Results

### Investigating tumor biology in a *Rb1*(-/-); *Trp53*(-/-); *Pten*(-/-), T cell infiltrated, prostate tumor model

To address the lack of neoantigens in tumor cells in mouse models, we developed a novel prostate cancer mouse model based on the NINJA mouse (*17*) (**Fig. 1A, B**). NINJA mice, on the C57BL/6 background, have a FLPo mediated central exon inversion to turn on neoantigen expression, specific for CD8 and CD4 T cells. The FLPo recombinases can be turned on by administration of Cre recombinase, tamoxifen, and doxycycline (**Fig. 1A**). This inducible system allows for precise temporal control over neoantigen presentation, and thus T cell infiltration into the tumor. To establish a physiologically relevant model, we generated from uninduced NINJA mice *Rb1*(-/-) *Tp53*(-/-) *Pten*(-/-) (TKO) mouse prostate organoids, modeling a constellation of loss-of-function mutations in tumor suppressors which frequently cooccur in CRPC patients (*6*) (**Fig. 1B, C**), using electroporation of Cas9-gRNA complex (**Fig. 1B**). We orthotopically transplanted the organoids into the prostates of immunocompetent C57BL/6J male mice (**Fig. 1B**), induced neoantigen expression four weeks post-transplant, and collected the prostates 6-weeks and 12-weeks post-transplant (**fig. S1E, F**). To understand the impact of castration and phenocopy androgen deprivation therapy, we castrated the mice at 8-weeks post-transplant. Western blot confirmed successful depletion of RB1, TP53, and PTEN proteins in organoids following CRISPR-Cas9 targeting (**Fig. 1C**) and the TKO organoids exhibited slithered morphology (**fig. S1A**) and increased resistance to AR degrader treatment compared to controls (**fig. S1B**).

**Figure 1.**
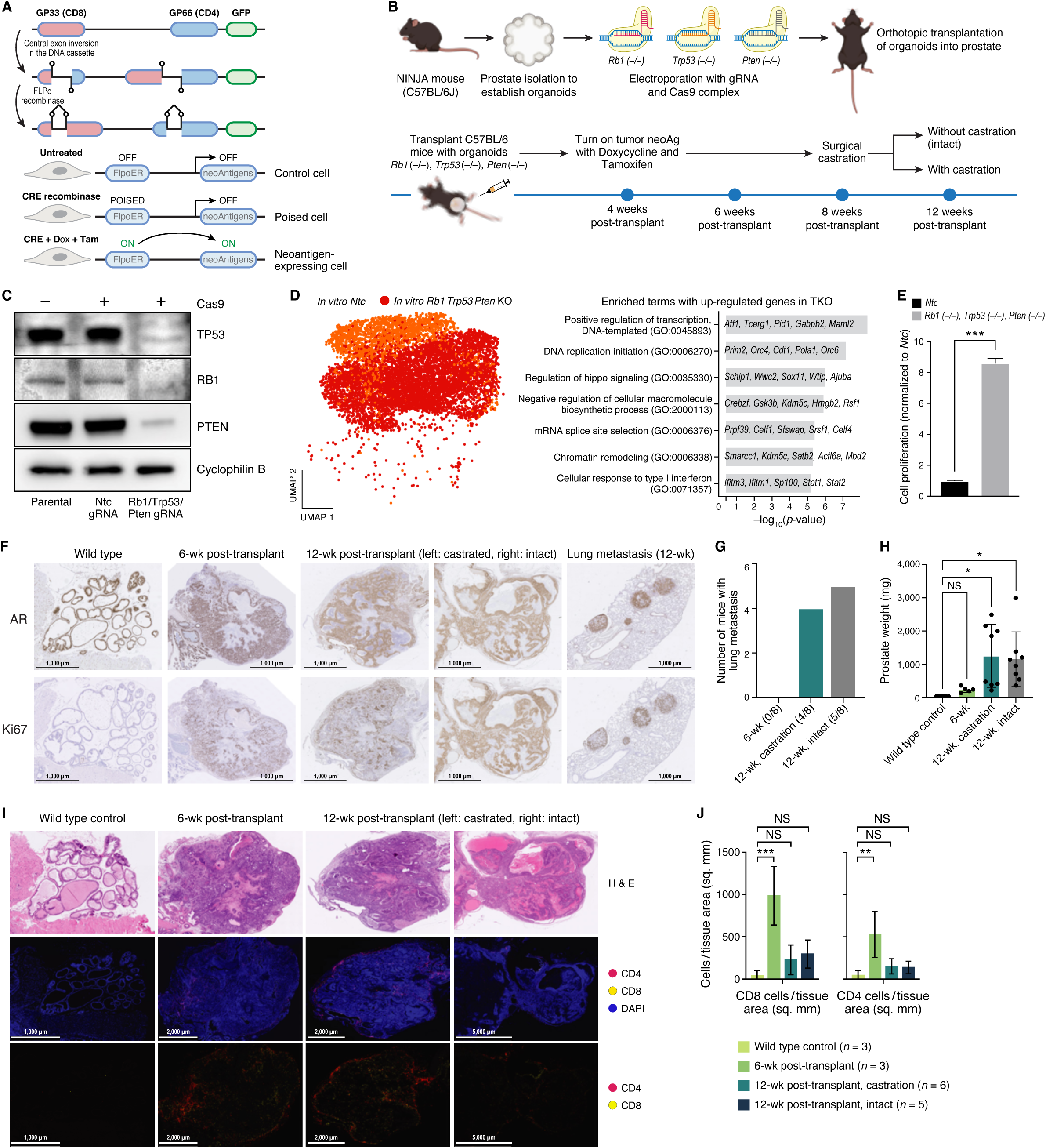
A prostate cancer tumor model with immune cell engagement. **A**, NINJA construct. Top: DNA construct. Bottom: Controlled neoantigen expression. After treatment CRE recombines, doxycycline (dox), and tamoxifen (tam), FLPo fused to estrogen receptor (ER) allows for nuclear localization, and neoantigen expression. **B,** Model generation. Top: Generation and transplantation of *Rb1* (-/-) *Trp53* (-/-) *Pten* (-/-) (TKO) mouse prostate organoids, with ribonucleoprotein (RNP) CRISPR/Cas9. Bottom: Experimental time course. **C,** Validated knockout of RB1, TP53, and PTEN. Western blot of protein levels in each condition (columns). Cyclophilin B: loading control. **D** Distinct cell profiles of *Rb1*^-/-^*Trp53*^-/-^*Pten*^-/-^ TKO mouse prostate organoids. Left: Uniform Manifold Approximation and Projection (UMAP) embedding of scRNA-seq profiles from non-targeting control (NTC, orange) and TKO (red) organoids. Right: Gene Ontology (GO) terms (y axis) enriched (-log_10_(P-value), for genes upregulated in TKO vs. NTC organoid cell profiles. **E,** Enhanced proliferation of TKO organoid cells. Mean *in vitro* proliferation (y axis) of cells from NTC and TKO organoids (x axis). Error bars: standard deviation (SD). *** P <0.001, Welch’s t test. **F-J,** Tumorigenesis by transplanted TKO organoids. F, Representative images of prostate or lung tissue after transplantation with NTC (WT) or TKO organoids (columns) stained by immunohistochemistry (IHC) for androgen receptor (AR) (top) and Ki67 (bottom). G, Number of mice (y axis) with lung metastasis at different time points post transplantation of TKO organoids (x axis). H, Mean prostate weight (y axis, mg) from WT and TKO transplants (x axis). Error bars: standard deviation. * P <0.05, one-way ANOVA. I, Representative images of prostate tissue from WT and TKO transplanted mice (columns) stained by hematoxylin and eosin (H&E, top), and DAPI, and imaged by immunofluorescence (middle and bottom) for CD8 (yellow), CD4 (red) and DAPI (blue). J, Density (y axis cells/mm^2^) of CD8 (left) and CD4 (right) T cells from WT and TKO transplants (x axis). NS: Not significant, * P< 0.05, **p < 0.01, ***p < 0.001; one-way ANOVA test.

ScRNA-seq of TKO and control (*Ntc*) organoids revealed upregulation of gene programs associated with positive regulation of transcription and DNA replication initiation, as well as of the type I interferon response (*Stat1* and *Stat2*), and of negative regulators of the Hippo pathway (*Wtip* and *Ajuba*) (**Fig. 1D and fig. S1C,D**), as early as ∼5 days post editing. This rapid *ex vivo* induction of distinct expression programs strongly suggests this to be a cell-autonomous and direct effect of RB1, TP53, and PTEN loss. This was also accompanied by significantly enhanced proliferative capacity (**Fig. 1E**), all consistent with the tumor suppressive role of RB1, TP53, and PTEN.

We characterized the model as castration resistant by high AR expression, metastasis and proliferation even with androgen depletion. Specifically, immunohistochemistry (IHC) of wild-type prostate tissue showed strong AR and negligible Ki67 staining, consistent with a non-malignant state (**Fig. 1F**), while by 6-weeks post-transplant, tumors developed high AR expression and high Ki67 staining across the entire tumor mass, demonstrating robust proliferation (**Fig. 1F**). At 12-weeks, tumors from both castrated and intact cohorts maintained strong AR and Ki67 staining (**Fig. 1F**). The high AR tumor phenotype was also retained in distant lung metastatic nodules at 12-weeks, in both castrated and intact mice (**Fig. 1F, G**). Given the equal prevalence of distant lung metastasis and increase in prostate weight **(Fig. 1H)** in both castrated and intact groups at 12-weeks, the TKO model can be classified as castration resistant, with high AR expression, where tumor cells are able to sustain AR signaling and proliferation despite androgen depletion. This underscores the aggressive nature of the AR-high cells in this model and its physiological relevance, given that reactivation of AR signaling is the most common driver of CRPC in patients (*2, 5*).

CD8^+^ and CD4^+^ T cell infiltration in the tumor parenchyma (as marked by an expert pathologist) after neoantigen induction showed dynamic spatial organization patterns over time based on a four-color immunofluorescence (IF) assay for CD8, CD4, EPCAM and DAPI (**Fig. 1I, J**). An initial high T cell infiltration at 6-weeks was reduced at 12-weeks, independent of castration (**Fig. 1I, J**). These T cells were largely neoantigen-specific, based on flow cytometry of MHC class I tetramer-positive CD8^+^ T cells (**fig. S1G, H**). These data support our system as a rapid, physiologically relevant novel prostate cancer model.

### Spatial transcriptomics reveals dynamic cellular reorganization during prostate tumor progression and metastasis

To comprehensively dissect the cellular composition, cell states and spatial organization of the prostate TME and metastatic sites over time, we performed high-resolution spatial transcriptomics of 5,100 genes, comprising 5,000 Xenium panel genes plus a custom 100-gene panel tailored to prostate biology (**Fig. 2A, fig. S2A-C**). The 100 gene-panel was designed through an interactive and iterative process between an autonomous AI agent (SpatialAgent (*19*)) and a human scientist (**Fig. 2A, table S1)**. We profiled two prostates each from wild-type control mice, 6- and 12- weeks post-transplantation with TKO organoids (with castration), as well as three lung tissues with multiple metastatic lesions (from castrated mice) (**Fig. 2A-C**).

**Figure 2.**
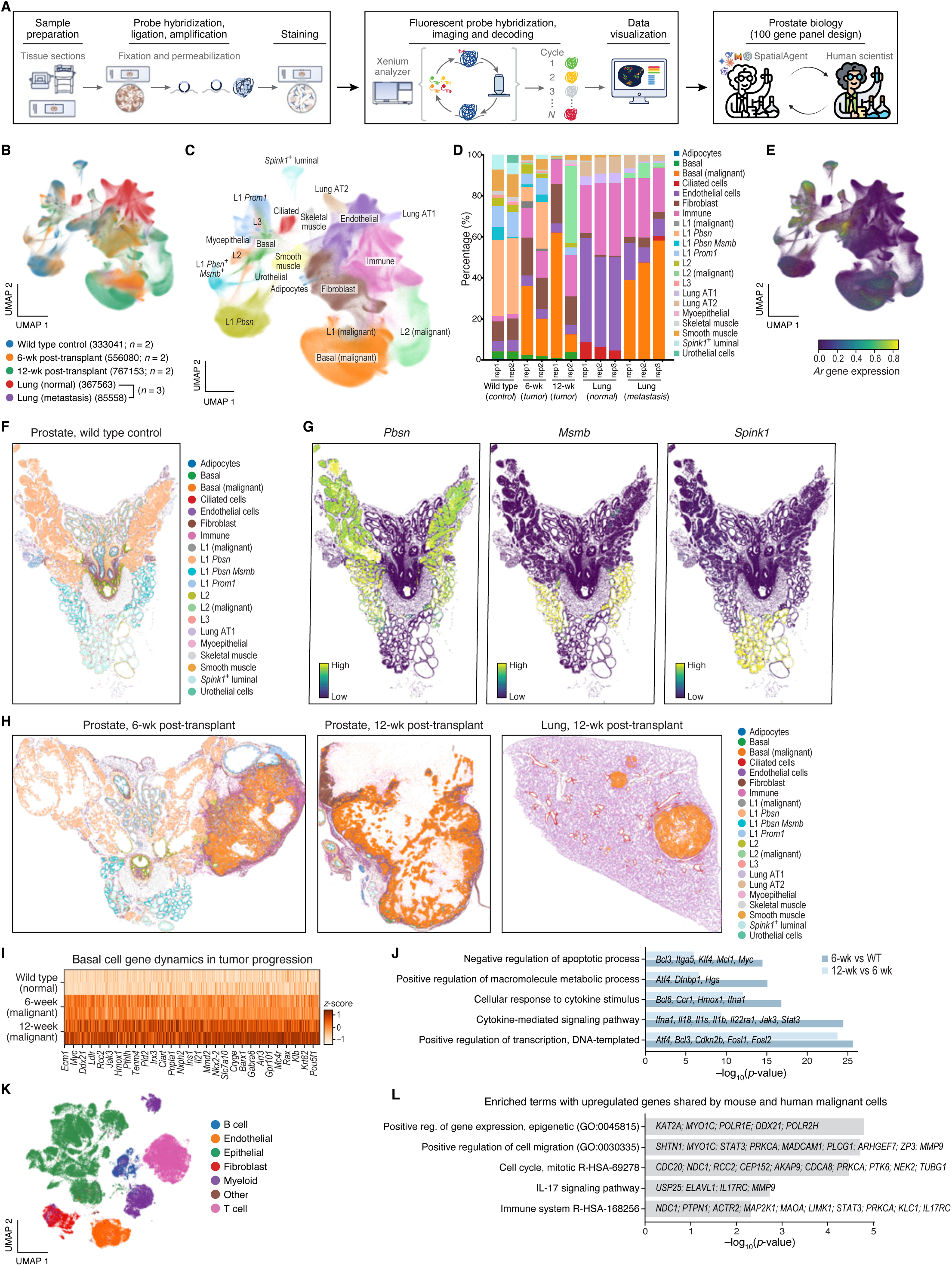
Malignant cell states and organization from primary prostate tumor to lung metastasis. **A,** Experimental workflow. **B-E** Changes in malignant and TME cell populations during tumorigenesis. UMAP embedding of segmented cell profiles from spatial transcriptomics (B,C,E) colored by sample (B), cell type (C) or AR gene expression (E). D, Cell type proportions (y axis) in tumor and normal portions of each section in all animals from each time point (x axis). **F-H,** Spatial cell type organization in primary tumors and lung metastases. Wild type control prostate (F,G), primary 6w (H, left) and 12w (H, center) tumors and lung metastases (H, right) colored by cell type annotations (F,H) or expression level of key marker genes (G) based on Xenium spatial trascriptomics. **I,J** Genes up-regulated in basal cells during tumorigenesis. **I**, Expression changes (Z-score) in basal cells from wild type, 6-weeks, and 12-weeks tumors (rows) for differentially expressed genes (P < 0.05, T-test on linear regression slope) (columns). **J,** Gene Ontology terms (y axis) enriched (-log_10_(p-value), x axis) in genes up-regulated (FDR<0.05) in mouse and human malignant cells. Select genes are highlighted. **K, L** Malignant cells in human prostate tumors express similar programs to mouse. K, UMAP embedding of scRNA-seq profiles from three prostate cancer patient studies (*23–25*) (n=26 patients) colored by cell type. L, Gene sets (y axis) enriched (-log_10_(p-value), x axis) for genes up-regulated (FDR<0.05) in malignant *vs*. non-malignant prostate epithelial cells from patient tumors.

Unsupervised clustering partitioned the 1,958,742 segmented cells from all samples (five tissue types) into 21 clusters, revealing substantial heterogeneity within the prostate epithelial cell compartment, fibroblasts, and immune cells (**Fig. 2B–D, fig. S2D**). In the wildtype mouse, there were three transcriptionally and spatially distinct subsets of L1 cells: a *Pbsn*-high population in the anterior lobe, a *Msmb*-high population in the lateral lobe and a *Spink1*-high population in the ventral lobe (**Fig. 2F, G**). AR was highly expressed in healthy prostate epithelial cells, malignant epithelial basal cells, and L2 cells (**Fig. 2E, fig. S2E**), consistent with the well-established role of AR signaling in prostate adenocarcinoma (*20*). Interestingly, AR expression was also high in adipocytes, smooth muscle and skeletal muscle cells (**Fig. 2C, E**).

In prostate tissue 6-weeks post-transplantation, early tumor foci emerged as nascent clusters of malignant cells, displacing normal prostate architecture (**Fig. 2H**, left, **fig. S2I**). By 12-weeks, primary tumors in castrated mice presented as large, dense masses predominantly composed of malignant cells with variable squamous differentiation, with significant distortion of the surrounding normal tissue (**Fig. 2H**, middle, **fig. S2J**). Keratin debris accumulated in the core of 12-week tumors from castrated mice (**Fig. 2H**, middle (white spaces)). The three lung tissues analyzed presented with multiple, relatively smaller, foci of metastatic lesions (**Fig. 2H**, left, **fig. S2K**).

Concomitantly, cell composition shifted dramatically along tumor progression (**Fig. 2D**). In primary tumors there was progressive and significant depletion of normal prostate L1, L2 and L3 cells, and an expansion of malignant-basal (in most tumors) and malignant-L2 (in one primary tumor and some metastases) *de novo* tumor cell states (**Fig. 2D, fig. S7A**). Interestingly, L2 cells (*Sca1*^+^ and *Psca*^+^) have been reported to contain a stem cell-like gene program that participates in the repair and regeneration of prostate (*22*). In the lung tissues containing metastatic foci, malignant prostate basal epithelial cells constituted a substantial proportion of the total cellularity of the lesion, confirming successful metastatic colonization, infiltration into the lung parenchyma and co-localization with host lung epithelial cells (**Fig. 2H, fig. S2K**).

The gene expression profiles of metastatic basal cells were very similar to those of malignant basal cells in prostate at 6- and 12-weeks, demonstrating the aggressive nature of primary malignant basal cells (**fig. S2F**). Basal cell profiles shifted along tumor progression, with a progressive increase in the expression of genes associated with inflammatory program (*Jak3* and *Stat3*), cytokine-mediated signaling (*Ifna1*, *Il18*), and negative regulation of apoptosis (*Bcl3*, *Mcl1*, *Myc*), key expression changes known to drive tumor progression (**Fig. 2I, J, fig. S2G**).

Comparing the expression profiles of malignant cells in our mouse model to those of human malignant epithelial cells from scRNA-seq profiles of 25 prostate cancer patients (**Fig. 2K, fig. S2L-N**) revealed a robust gene signature shared by malignant basal cells across the two species (**Fig. 2L, table S2**) (*23–25*). The conserved program is enriched for genes from key cancer processes, like the cell cycle, and specific pathways like immune system regulation and the IL-17 signaling pathway (**Fig. 2L**). This shared inflammatory module was enriched for key JAK-STAT signaling effectors—most notably *Stat3*—as well as downstream mediators of invasion and inflammation such as *Mmp9* and *Il17rc* (**Fig. 2L**). Collectively, our analysis provides a spatial and molecular map along trajectory of prostate tumor evolution in a physiologically relevant prostate tumor model, characterized by progressive loss of normal glandular architecture, emergence of transcriptionally distinct malignant states, and reprogramming of basal and (in some tumors) L2 epithelial cells.

### Immune-stromal niche remodeling and niche-CD8 T cell communication along tumor progression

The prostate cancer TME has been reported to be highly immunosuppressive, with multiple mechanisms hindering the effectiveness of CD8 T cell-mediated anti-tumor immunity (*26*). In the immune compartment, it is known to recruit myeloid-derived suppressor cells (MDSCs), neutrophils, regulatory T cells, and tumor-associated macrophages, and each can influence T cell differentiation, in prostate and other cancer types (*27, 28*). The stroma also functions as a critical immunomodulatory barrier in prostate and other cancers. Its dense collagen and matrix deposition physically sequester T-cells within the peritumoral space as well as actively drive their differentiation toward dysfunctional phenotype (*29, 30*).

Given the established role of CAFs in tumor progression in multiple cancers (*31, 32*), and the profound disruption of normal prostate architecture in our model, we first examined the changes in normal fibroblasts and cancer associated fibroblasts in our tissues (**Fig. 3A, B**). In wild-type prostate, the stromal compartment was dominated by quiescent *Pi16*⁺ and *Col231*⁺ fibroblasts (**Fig. 3A-C, fig. S3A, B**), whereas in 6-week tumors, normal fibroblasts were largely replaced by a distinct *Ccl2-Jak2* double positive inflammatory fibroblast population (**Fig. 3A-C, fig. S3C)**. While normal *Pi16⁺* fibroblasts were excluded from the tumor parenchyma, *Ccl2-Jak2*⁺ inflammatory CAFs were primarily tumor-adjacent and infiltrated, including in the tumor core (**Fig. 3C**). At 12-weeks, this was further augmented by the emergence of *Mest*^+^ and *Fosl1*^+^ CAFs that were tumor infiltrated, and further reduction of quiescent ones (**Fig. 3A-C, fig. S3I**). A similar shift is observed in lung metastases (**Fig. 3A, fig. S3K**). Notably, previous work suggests that CCL2 expression in the TME plays a role in promoting cancer cell proliferation and migration (*33–35*). Fibroblasts’ heterogeneity in the TKO mouse model was quite concordant with that in human prostate cancer data (**Fig. 3G, fig. S3L, M**). In both human and mouse tumors, the Ccl2+Jak2+ inflammatory CAFs dominated, followed by Col23a1+ and Pi16_+_ normal fibroblasts (**Fig. 3G**). This striking similarity in the proportions of major fibroblast populations supports the physiological relevance of our TKO mouse model.

**Figure 3.**
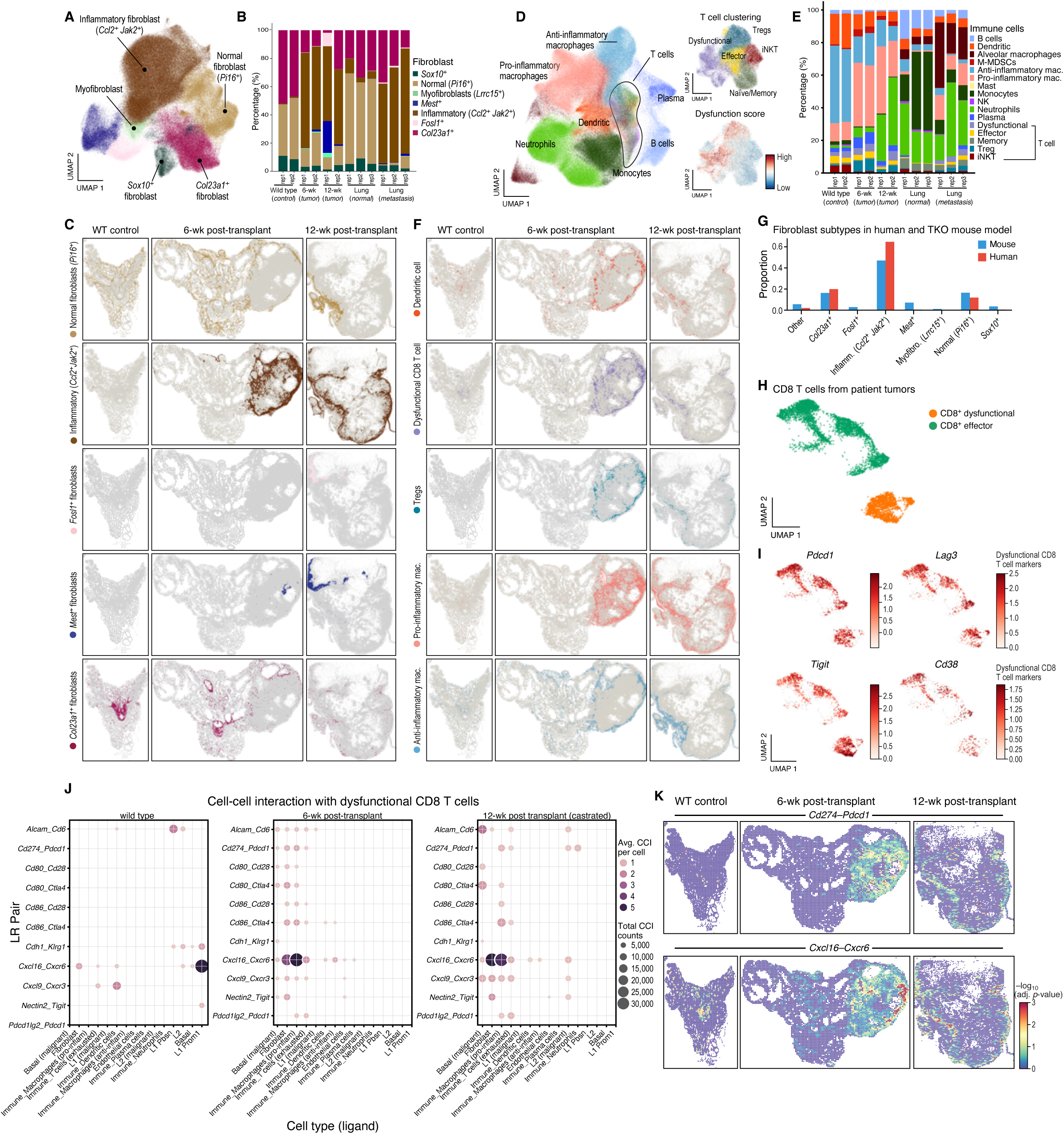
Immune-stromal niche remodeling and communication during tumor progression. **A-C,** Inflammatory *Ccl2-Jak2*^+^ CAFs rise in prostate tumors and metastases. A, UMAP embedding of RNA profiles of segmented fibroblasts from spatial transcriptomics of wild type prostate, and normal and tumor prostate tissue at 6-and 12-weeks post-transplantation colored by cell type annotation. B, Proportion (y axis) of segmented cell profiles from each fibroblast subset in each tissue region, from each animal (x axis). C, Spatial transcriptomics sections from each animal (columns) colored by each fibroblast subset (rows). **D-F,** Inflammatory macrophages and neutrophils rise in prostate tumors and metastases. D, UMAP embedding of RNA profiles of segmented immune cells from spatial transcriptomics of wild type prostate, and normal and tumor prostate tissue at 6-and 12-weeks post-transplantation colored by cell type annotation. E, Proportion (y axis) of segmented cell profiles from each immune cell subset in each tissue region, from each animal (x axis). F, Spatial transcriptomics sections from each condition (columns) colored by selected immune cell subsets (rows). **G-I**, Shared TME features between mouse and human prostate tumors. G, Proportion (y axis) of cells from different fibroblast subsets (x axis) in the mouse model (blue) and scRNA-seq of human prostate tumors (red; data as in Fig. 2K). H,I, UMAP embedding of scRNA-seq profiles of CD8 T cells from human prostate tumors (data as in Fig. 2K) colored by state (H) or by expression of key immune checkpoint and activation markers (I). **J,K,** Inflammatory fibroblasts and macrophages are key signaling partners to dysfunctional CD8^+^ T cells. J. Total number of cell-cell interaction (CCI) (dot size) and average number per cell (dot color) for receptor-ligand pairs (rows) between receptors (second gene name) expressed on dysfunctional CD8 T cells and ligands or co-receptors (first gene name) expressed by various proximal (within 30 μm) cell types (columns) in wild type prostate (left), 6 week (center) or 12 week (right) prostate tumors (tumor segment of section). K, Spatial transcriptomics sections of wt prostate (left), 6 week (middle) and 12 week (right) prostate tissue colored by significance (-log10(FDR), permutation test (*42*)) of each receptor-ligand pair.

Myeloid cell composition and spatial organization were also drastically remodeled in the prostate tumors *vs*. healthy prostate, as well as in lung metastases (**Fig. 3D, E, fig. S3D-F**). In healthy wild type prostate, dendritic cells and anti-inflammatory macrophages (*Mrc1*^+^*Cd163*^+^*Lyve1*^+^ (*36, 37*)) were the predominant cell populations, while their proportions were reduced in tumor bearing conditions, where pro-inflammatory macrophages (*Spp1*^+^*Fn1^+^Cxcl16^+^*(*36, 37*)) and neutrophils expanded by 6 and 12-weeks (with castration), respectively (**Fig. 3D-F, fig. S3J**). Moreover, pro-inflammatory macrophages were infiltrated in the tumor parenchyma, directly surrounding the tumor, whereas anti-inflammatory macrophages and dendritic cells were excluded from the tumor core (**Fig. 3F, fig. S3J)**. In lung metastases, monocytes were decreased *vs*. normal lung, while pro-inflammatory and alveolar macrophages were highly elevated (**Fig. 3E, fig. S3K**). Anti-inflammatory and alveolar macrophages were excluded from the tumor parenchyma while pro-inflammatory macrophages were tumor infiltrated (**fig. S3K**). Interestingly, alveolar macrophages have been reported to play a role in premetastatic niche formation and suppressing antitumor T cell responses (*38, 39*).

There was an increase in the number of both dysfunctional and effector CD8 T cells in 6-week and 12-week tumors, compared to wild type control or the normal tissue at 6 and 12-weeks, roughly proportionally to the growth of the overall immune compartment in the tumor vs. the non-tumor tissue (**fig. S3G,H, Fig. 2D, 3E**). Dysfunctional CD8 T cells were both tumor infiltrated and on its periphery (**Fig. 3F, fig. S3J**). Regulatory T cells (T_regs_) were prevalent in tumor tissue and absent in normal or histologically defined non-malignant tissue (**Fig. 3D-F, fig. S3J**). In one 12 week tumor, both dysfunctional CD8 T cells and Treg were more excluded at 12 weeks (**Fig. 3F, fig. S3J**). Comparing the CD8 T cell phenotypes in the mouse model to intratumoral CD8 T cells from 25 patients with prostate cancer showed that human prostate intratumoral CD8 T cells segregate into two effector-like and dysfunctional-like cells (**Fig. 3H, Methods**), both expressing coinhibitory receptors like *Pdcd1*, *Tigit*, *Lag3*, and *Cd38* (**Fig. 3I**), consistent with the up-regulation of inhibitory molecules in T cell activation (*40, 41*), and analogous to the states observed in the mouse model, supporting its relevance.

Spatially constrained ligand-receptor (LR) interaction analysis between dysfunctional CD8 T cells and all other cell types proximal to them (within 30 μm of the segmented cell (*37, 42*)) in the tumor niche revealed that tumor progression was accompanied by increasing complexity of potential signaling into exhausted CD8 T cells (**Fig. 3J**), mostly from Ccl2+ fibroblasts and inflammatory macrophages, and by 12 weeks also increasingly from malignant cells. Among the most enriched LR pairs were *Cxcl16*–*Cxcr6, Cd274*–*Pdcd1* (PDL1–PD-1), and *Cd80*-*Ctla4*, indicating activation of immune checkpoint pathways and chemokine-driven immune modulation (**Fig. 3J, K**). Together, these data demonstrate that prostate cancer progression in our model is accompanied by profound spatial reorganization of the immune and stromal microenvironment, resulting in the emergence of a highly inflammatory and stromal niche that may facilitate immune evasion.

### Zman-seq charts the temporal dynamics of infiltrating immune cells

To understand how infiltrating immune cells arriving from the circulation, including T cells, monocytes and neutrophils (**Fig. 3D,E**) are remodeled and reprogrammed by the prostate cancer TME, we next used Zman-seq (*18*) (**Fig. 4A**). Briefly, in Zman-Seq, immune cells in peripheral blood of one mouse are labeled *in vivo* along a time course using anti-CD45 antibodies labeled with distinct fluorescent dyes (one per time point), and this “time-stamps” each cell at the point of labeling. If timestamped cells migrate into tissues (here, the prostate tumor), those that infiltrate later will carry different dyes than those that infiltrated earlier. At the end point, we dissociate the tumor, sort the different immune cells with FACS by their district dyes, and then profile each labeled population by scRNA-seq (*18*).

**Figure 4.**
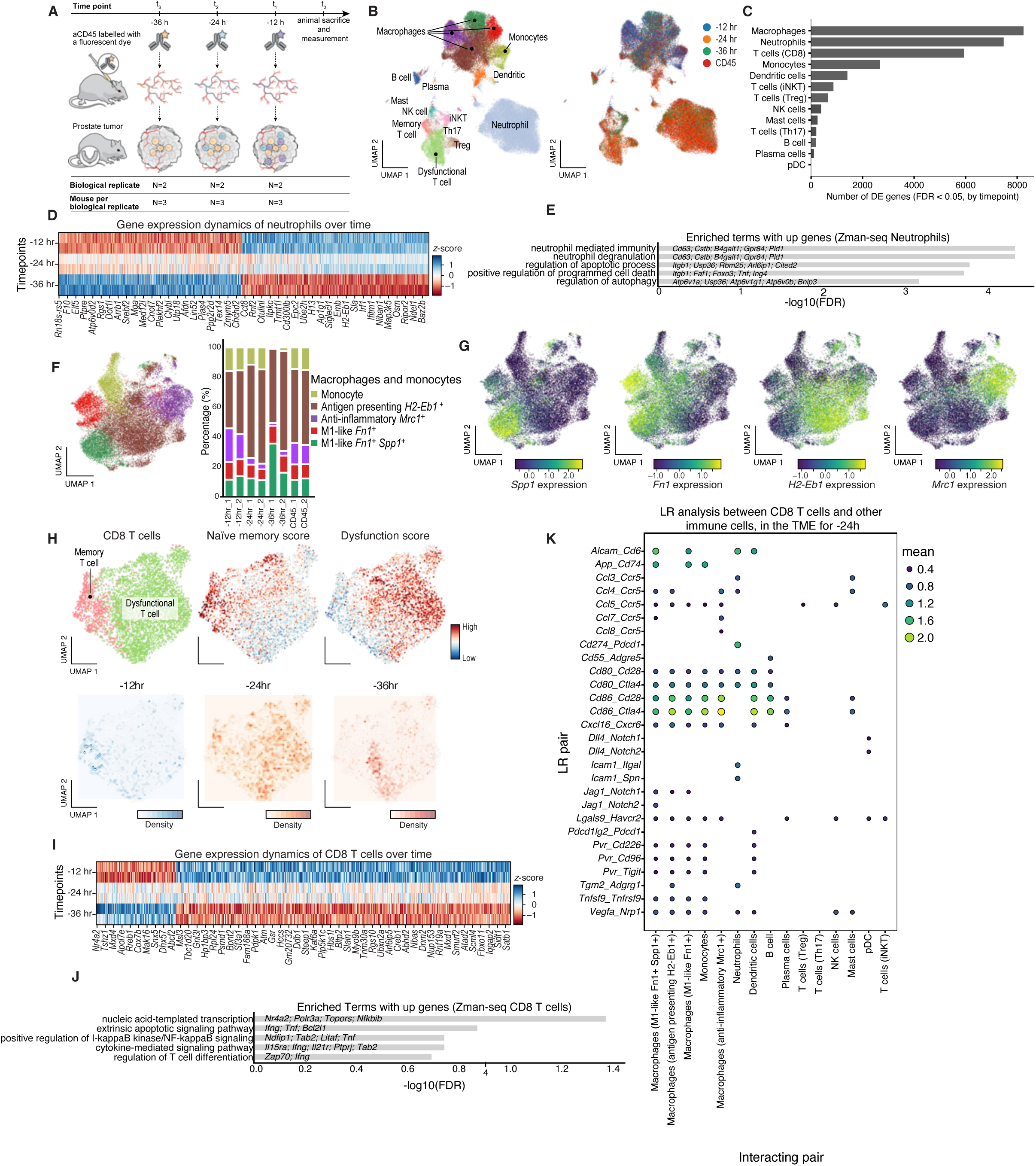
Reprograming of infiltrating immune cells by the TME revealed by Zman-seq. **A**, Zman-seq experiment. **B,C** Macrophage, CD8 T cells and neutrophil profiles shift by TME exposure. B, UMAP embedding of all CD45^+^ immune cell profile from Zman-seq colored by cell type (left) or duration in tumor (right). CD45 – all immune cells present in the tumor (labeled and unlabeled). Data from two biological replicates was integrated for each time point. C, Number of genes (x axis) differentially expressed between labeling times (FDR<0.05, t-test with overestimation of variance) in each immune cell subset (y axis). **D,E,** Reprogramming of infiltrating neutrophils by the TME. D, Mean normalized expression (z-score) for genes (columns) that are differentially expressed (*P*<0.05, linear regression, Wald test) in neutrophils between time points (rows). E, Gene Ontology sets (y axis) enriched (-log_10_(FDR), x axis) for genes differentially expressed in neutrophils with time. Key gene names are marked. **F,G,** Reprogramming of infiltrating macrophages by the TME. UMAP embedding of macrophage Zman-Seq profiles colored by macrophage subset (F, left) or key macrophage marker genes (G) and proportion (y axis) of macrophage subset at different time points following infiltration and the tumor overall (CD45) (x axis) (f right). **H-J,** Shift in T cell states following infiltration. H, UMAP embedding of CD8 T cells colored by subset (top left), naïve memory (top middle) or dysfunction (top right) scores, and by density of cells from each duration in the tumor (bottom). I, Mean normalized expression (z-score) for genes (columns) that are differentially expressed (P-value < 0.05, linear regression with pseudo-bulk, Wald test) in CD8 T cells between time points (rows). J, Gene Ontology sets (y axis) enriched (-log_10_(FDR), x axis) for genes differentially expressed in neutrophils with time. **K,** Prominent cell-cell interactions with CD8 T cells and other immune cells, that have been in prostate TME for 24 hours. Mean expression (dot size and color) of the interacting molecules in the interacting cells of ligand-receptor pairs (rows) between CD8 T cells and other immune cells (columns) at 24 hours of exposure. Red: interactions highlighted in the main text.

We applied Zman-seq to mice with 6-week tumors, the time of maximal immune infiltration (**Fig. 1I**, **Fig. 3**), by injecting anti-CD45 antibodies with different labels at -36h, -24h, and -12h prior to animal sacrifice (t=0) at 6-weeks (**Fig. 4A,B, fig. S4A-D**). We performed two replicate experiments, each with three labeled mice, with immune cells from the three mice (three prostate tumors) pooled for profiling. We annotated cells by clustering all CD45^+^ cells into to 17 subsets, spanning various lymphoid (T cells, B cells, NK cells, iNKT) and myeloid (macrophages, monocytes, dendritic cells, neutrophils) subsets (**Fig. 4B, fig. S4A-D,G**). For each subset, we identified genes differentially expressed at each timepoint *vs*. all other timepoints (**Fig. 4C**, FDR < 0.05, t-test with overestimations of variance). Macrophages, monocytes, neutrophils, and CD8 T cells had the most prominent changes in gene expression by TME exposure (**Fig. 4C**).

Infiltrating neutrophils progressively turned on an inflammatory gene program (**Fig. 4D,E**), a neutrophil-mediated immunity program and a neutrophil degranulation gene program, with greater expression of *Cd63*, *Cstb*, *B4galt1*, *Gpr84* and *Pld1,* as a function of time spent in TME. (**Fig. 4E**). Interestingly, this is the canonical neutrophil program that turns on in the early stages of wound healing; however, if the program does not turn off after an initial burst, then it leads to chronic inflammation and significant tissue damage (*43, 44*). In the context of a tumor, expression of an inflammatory program in neutrophils has been shown to promote tumor growth, metastasis, and immune evasion (*45, 46*).

Monocytes, which are often the precursors of tumor associated macrophages (*47, 48*), were present in higher proportion after shorter labeling times (-12h) but had drastically decreased with tumor residency time (-36h), supporting prostate TME mediated differentiation into macrophage cell fates (**Fig. 4F**). Similarly, MRC1^+^ anti-inflammatory macrophages steadily decreased in proportion with tumor residency (**Fig. 4F, G**), whereas FN1^+^SPP^+^ pro-inflammatory macrophages increased (**Fig. 4F, G**).

Among T cells, T_reg_ profiles showed an increase over time in the tumor in anti-stress (*Crebbp*, *Hsf1*), anti-apoptosis (*Casp3*, *Nr4a2*), and pro-cell cycle (*Ccnd2*, *Pkn2*) programs (**fig. S4E, F**), which may be adaptative to stress in the TME. In CD8 T cells, there is a progressive increase in the dysfunction score with longer tumor residency, concomitant with a decrease in the naïve memory score and in genes such as *Ccr7*, *Cd45ra* and *Cd95* (Fas) (**Fig. 4H**). Up-regulation of the dysfunctional programs occurs as early as 24h in the TME (**Fig. 4H**). As tumor residency increases (from -12h to -36h labeling), CD8 T cells shift to a state that is high in transcription and differentiation programs (**Fig. 4I,J**). Among the key genes up-regulated over time by the TME in tumor-infiltrating CD8 T cells are canonical effector cytokines *Ifng* and *Tnf*, the pro-survival factor *Bcl2l1* and the checkpoints *Pd1* and *Havcr2* (TIM3), consistent with activation followed by co-inhibition (**Fig. 4I,J**), a hallmark of the transition to a dysfunctional state (*49–51*).

To identify potential drivers of this transition in CD8 T cells, we performed a comprehensive ligand-receptor interaction analysis between CD8 T cells and other immune cells at 24 hours post entry (**Fig. 4K**). This analysis highlighted numerous key interactions, for example, *Pdcd1* (PD-1) on CD8 T cells interacting with *Cd274* (PD-L1) expressed by neutrophils, and *Ctla4* on CD8 T cells interacting with *Cd80* and *Cd86* on tumor associated macrophages and dendritic cells (**Fig. 4K**). Additional interesting interactions of *Lgals9* (Galectin-9) with *Havcr2* (TIM-3) and *Pvr* (CD155) with *Tigit* (**Fig. 4K**) further underscored the complex immunosuppressive network targeting CD8 T cells. These findings delineate the temporal onset of T cell dysfunction within the prostate tumor.

### Combined treatment with anti-PD1 and JAK1/2 inhibitor remodels the inflammatory niche and reduces tumor growth

The prostate TKO NINJA model shows dysfunction and progressive exclusion of tumor-engaged CD8 T cells, tumor infiltration of inflammatory cells (pro-inflammatory macrophages, *Ccl2*^+^*Jak2*^+^ CAFs, neutrophils) and elevated JAK-STAT and inflammatory signaling in basal malignant cells. Moreover, LR analysis of both spatial transcriptomics and Zman-seq data highlighted putative interactions between the inflammatory niche and CD8 T cell dysfunction. We therefore hypothesized that breaking inflammatory signaling in the niche could sensitize prostate tumors to ICB. Notably, human *Ccl2*^+^ CAFs also scored highly for the IL6-JAK-STAT3 signature overall (though individual genes vary in their expression between human and mouse), and JAK inhibition has recently shown promise in enhancing ICB in lung cancer and lymphoma by modulating T cell plasticity and myeloid programming. However, its impact and efficacy in prostate TME remains unexplored (*52, 53*).

To test this, we treated TKO tumor-bearing mice with anti-GP120, anti-PD1, JAK1/2 inhibitor (JAKi), and the combination of anti-PD1 + JAKi (**Fig. 5A**). The tumors were completely resistant to anti-PD1 monotherapy, as expected (*12, 13*), while JAKi monotherapy was non-significant in reducing tumor weight (**Fig. 5B)** but yielded a significant effect on tumor volume (**Fig. 5C, fig S5A,B**; eGaIT and eDOT contrast, **Methods**). However, the combination of anti-PD1 and JAKi significantly suppressed tumor volume growth and reduced tumor weight (**Fig. 5B,C, fig. S5A,B;** eGaIT and eDOT contrast, **Methods**). Histopathological analysis of tumor tissue highlighted loss of intact tumor epithelium and disorder of the remaining epithelium in the combination group relative to other three groups (**fig. S5C**). To characterize how the tumor’s architecture may change under therapy, we performed spatial transcriptomics (Xenium) on three tumors (from three different mice) from each of the four treatment arms using the ∼5,100 gene panel used earlier (**Fig. 5D,E, fig. S5C**).

**Figure 5.**
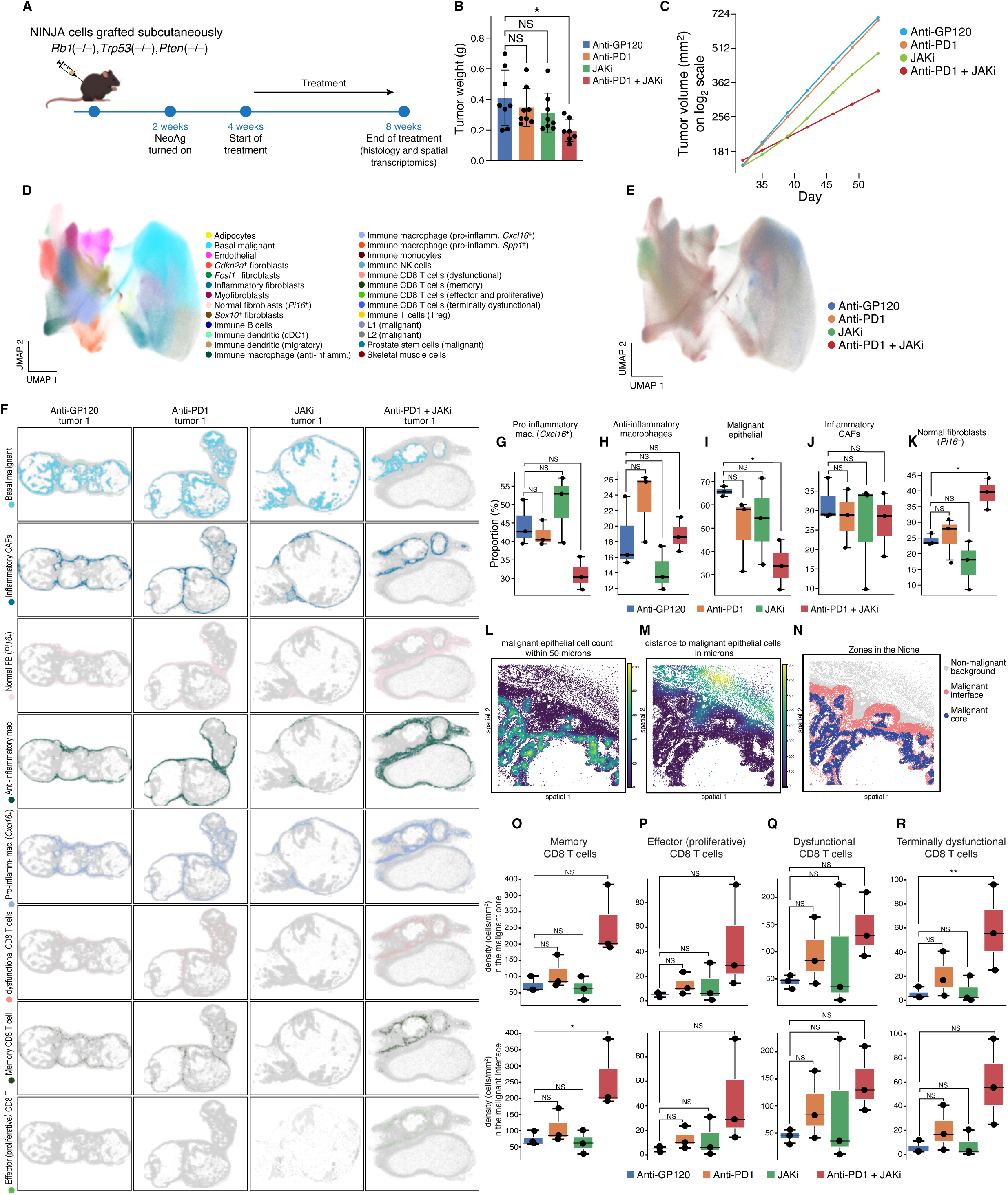
Combined anti-PD1 and JAK inhibitor treatment inhibits tumor growth and reshapes the TME. **A,** Experimental design. **B-C** aPD1 and JAKi combination inhibits tumor growth. B, Mean tumor weight (y axis, g) comparison at the 8-week end point for each regimen (colors). *p < 0.05; NS-nonsignificant, One-way ANOVA test. Error bars: standard deviation (SD). C, Tumor volume (y axis, mm^3^) over time (x axis) for each regimen. **D-R,** Shift of tumor and TME cell composition and organization under combination treatment. D,E, UMAP embedding of segmented cell profiles from Xenium spatial transcriptomics (dots) from all conditions (12 tumors) colored by cell subset annotation (D) or regimen (E). F, A representative spatial transcriptomics tumor section from each condition (columns) colored by key cell subsets (rows). **G-K**, Proportion (y axis) of key cell subsets (label on top) in each regimen (x axis). Box plots: Center line – median, box limits – upper and lower quartiles, individual data points - independent biological replicates (n=3 mice per group). *p < 0.05; NS-nonsignificant, one-way ANOVA test. **L-R**, T cell density changes in distinct regions following treatment. **L-N**, Region delineation. Example tumor region in a section, colored by epithelial cell number within 50μm (L), distance to malignant epithelial cells (M) and region label classification (**Methods**) (N). **O-R**, Density, (cells/mm^2^, y-axis) in the malignant core (top) and interface (bottom) of key cell types (label on top) in each treatment (x axis). Box plots: Center line – median, box limits – upper and lower quartiles, individual data points – independent biological replicates (n=3 mice per group). *p < 0.05; **p < 0.01; NS-nonsignificant, one-way ANOVA paired with Tukey’s Honestly Significant Difference (HSD) test.

Analysis of individual cell profiles in the spatial transcriptomic data revealed treatment-induced remodeling of the cell composition in the tumor niche by the combination therapy (**Fig. 5D-K**). Mirroring the reduced tumor volume and weight, the combination of anti-PD1 and JAKi significantly reduced the proportion of malignant epithelial cells in the tumor (**Fig. 5I,** p =0.049). While anti-inflammatory macrophages did not show net change with combination therapy (**Fig. 5H**), the combination was marginally synergistic in reducing the proportion of pro-inflammatory macrophages compared to the individual effects (beta coefficient of -16.193, p-value=0.054, **Fig. 5G**, **Methods**). Although the combination did not change inflammatory CAF proportions in the tumor (**Fig. 5J**), it significantly increased normal (*Pi16*^+^) fibroblasts (**Fig. 5K**, p-value = 0.033), in a highly synergistic manner (beta coefficient = 22.222, p-value = 0.013, **Methods**), compared to the individual effects of anti-PD1 (unchanged) and JAKi (nominally (insignificantly) decreased). Therapy-induced emergence of normal/quiescent fibroblasts has been shown to be tumor regressive and therapeutically beneficial in cancer models (*54, 55*).

To quantify T-cell infiltration into distinct tumor compartments of the TME as a function of treatment, we developed a multi-step spatial analysis pipeline (**Fig. 5L-R; Methods**) to define malignant core, malignant interface and normal background regions in the tumor (**Fig. 5L-N, Methods)** (*56, 57*). Interestingly, combination treatment was associated with a significant increase in the density of memory CD8 T cells in the malignant interface region (**Fig. 5O,** p-value = 0.030), while the combination did not significantly change the density of effector (proliferative) and dysfunctional CD8 T cells between the tumor core and interface regions (**Fig. 5P,Q)**. However, the density of terminally dysfunctional CD8 T cells was significantly increased in the malignant core (p-value = 0.0046, **Fig. 5R**). Our observation of reduced tumor growth alongside an increased frequency of terminal dysfunctional CD8+ T cells is consistent with the ’proliferative burst’ model, where immune checkpoint blockade triggers the differentiation of effector and progenitor dysfunctional cells into more cytotoxic but terminal dysfunctional state (*58, 59*). Thus, our data add to growing evidence that the efficacy of checkpoint inhibition may be enhanced through rational combination strategies informed by mechanistic understanding of the disease, in a physiologically relevant tumor model.

## Discussion

By combining spatial transcriptomics (Xenium) with Zman-seq, our work provides insight into the functional architecture of the prostate TME in an immunocompetent mouse model, enabling the direct visualization of tumor-niche interactions, and provides the first *in vivo* temporal map of immune cell reprogramming within a prostate tumor.

Our analysis characterized the changes in the prostate TME over weeks as the tumor establishes, grows and metastasizes, and over days as immune cells migrate from the periphery and infiltrate. We propose a model where macrophages, CAFs, and transformed basal cells form an inflammatory triad in the tumor niche and engage in paracrine signaling that promotes CD8 T cell dysfunction or exclusion. Expression analysis revealed activation of interferon and JAK/STAT signaling pathways in tumor cells, inflammatory macrophages and inflammatory CAFs.

Short term upregulation of inflammation is critical for generating an anti-tumor response and wound repair, however, chronic inflammatory signaling in a tumor, that some consider as a “wound that does not heal”, leads to tumor growth, metastasis, and immune evasion (*60–62*). Our findings suggest that the inflammatory triad in the TME establishes a tolerogenic niche that drives T cell exhaustion, even in the presence of neoantigens and infiltrating CD8 T cells. Notably, the NINJA TKO model compared well in terms of cell composition and states to human prostate tumor biology. In particular, the inflammatory wiring of malignant cells, the specific enrichment of iCAFs, and the distinct phenotype of dysfunctional CD8 T cells are all conserved between murine and patient data.

Based on these findings and recent studies in other tumors (*52, 53*), we demonstrate a potential therapeutic synergy of PD-1 blockade and JAK inhibition in our animal model, and its association with a multi-pronged reprogramming of the tumor niche. In addition to reduction in malignant cells, combination treatment led to a significant increase in normal (*Pi16*^+^) fibroblasts in the niche overall and a significant increase in the density of dysfunctional and memory CD8 T cells in the malignant core or interface region, respectively. This suggests that the combination treatment not only debulks the tumor but also fosters a more immunologically active and ’normalized’ niche (*63, 64*).

The use of multiplex CRISPR-Cas9 genome editing in organoids to generate our model highlights the scalable nature of the platform for interrogating genotype-immune phenotype relationships in prostate cancer and is one that is flexible to modeling other common PCa alterations, like Foxa1 and c-Myc overexpression with lentivirus addition. In this sense, our model is also a flexible platform for translational immuno-oncology. One remaining limitation is that the induced model neoantigens used, while powerful, may not be identical to the spectrum and kinetics of antigen evolution in human disease.

In sum, this study exemplifies how understanding tumor biology with novel models and spatial-temporal-omics technologies can uncover niche-driven resistance mechanisms and nominate rational combination strategies.

## Methods

### Animal experiments

All mice were housed and maintained at Genentech in accordance with American Association of Laboratory Animal Care guidelines. All animal studies were conducted under the approval of the Institutional Animal Care and Use Committees of Genentech Lab Animal Research and were performed in an Association for the Assessment and Accreditation of Laboratory Animal Care (AAALAC)-accredited facility. C57/BL6 and NOD SCID Gamma (NSG) mice were purchased from the Jackson Laboratory.

For orthotopic transplants, 8-12 weeks old C57/BL6 mice were purchased from Jackson laboratories, and 10^6^ cells were injected into the anterior lobe of the prostate. Cells were prepared after dissociating 3D organoids into single cell suspension. For orthotopic transplantation into each mouse, 10^6^ cells were resuspended in 10μl of Matrigel and 10μl of organoid media, a total volume of 20μl. For sub-cutaneous injections, organoids were dissociated, and 10^6^ cells were resuspended in 50μl of Matrigel and 50μl of organoid media, total volume of 100μl. This suspension was injected in the right flank of 8-12 weeks old C57/BL6 or NSG mice.

### Surgical castration

Surgery was performed in accordance with the IACUC’s Rodent Survival Surgery Guidelines. Animals were anesthetized with isoflurane. The abdomen was shaved, and the skin was disinfected with three times betadine or chloraprep swabs and isopropyl alcohol. Eye ointment or lubricant was applied to both eyes. The skin of the lower abdomen was incised longitudinally with a sterile scalpel. The peritoneum was incised, approximately 0.5-1 cm incision. Blunt forceps were used to pull out the testis, vas deferens, and epididymis. The attached fat was pulled to expose testis, vas deferens and epididymis. Vas deferens, epididymis, and testicular arteries were cauterized. Appropriately sized absorbable sutures were used to close the peritoneum. Sterile, appropriately sized wound clips were used to close the skin incision.

### Generation and culture of mouse prostate organoids

All cell lines and organoids were periodically tested and found negative for a mycoplasma test (Lonza #LT07-318). Cell lines used in this study were maintained in a 37°C and 5% CO_2_ incubator. Murine organoids were established from adult mouse prostate taken from a C57/BL6 mouse, as previously described (*65*). Briefly, the entire mouse prostate was harvested, and physically dissociated into smaller pieces, with sterile razor blades. Next, it was digested with collagenase type II (Gibco) for about 1hr at 37°C with gentle shaking. Next, TrypLE (Gibco) was added for digestion at 37°C until a single-cell suspension was obtained. Organoids were washed 3 times with cold PBS after TrypLE digestion. Organoid media was prepared as previously described (*65*), supplemented with Y-27632 (10 μM) to inhibit anoikis. Finally, cells were filtered through FACS tubes and seeded for organoid culture. Bulk isolated prostatic epithelial cells were embedded in 35μl drops of basement membrane extract (Matrigel, Corning) and overlaid with mouse prostate organoid medium.

### Electroporation of Cas9-sgRNA ribonucleoprotein (cRNP) complex

Nucleofection was performed using a basic kit for primary mammalian epithelial cells (Lonza,VPI-1005) or kit R (Lonza, VVCA-1001) for prostate organoids or a home-made nucleofection solution (150 mM KH2PO4, 24 mM NaHCO3, 3.7 mM glucose, 7 mM ATP (adenosine triphosphate)-disodium salt, 12 mM MgCl2, pH 7.4). 500,000 dissociated organoid cells were resuspended with nucleofection buffer, cRNP complexes, and electroporation enhancer (IDT 1075915, 1:1 molar ratio to cRNP) in a total volume of 100μL. The cell suspension was transferred to a nucleofection cuvette and nucleofected using Lonza Amaxa Nucleofector II, program T-030 for prostate cells. Cells were centrifuged and seeded for culture as organoids) in Matrigel, as mentioned earlier.

### Growth assay

For organoid growth assays, organoids were resuspended in PBS and CellTiter-Glo mix (Promega G7570), in a 1:1 ratio. Three to five replicates were performed for each data point. Experiments were repeated three independent times. All values were normalized to a reference sample in the experiment.

### Incucyte assay

Organoid formation and growth were analyzed using the Organoid QC module on an Incucyte SX5 Live-Cell Analysis System (Sartorius). Organoids suspended in a 10μl matrigel dome were carefully seeded in the well center of Corning 48-well plates. Following a 15min incubation at 37°C for matrigel polymerization, 250μl of media was added to each well, and the plate placed in the Incucyte. Brightfield images were captured every 4 hours for 7 days using a 4x objective, then segmented and analyzed with the integrated Incucyte Organoid software module. Total organoid area per well and average eccentricity readouts were used to quantify organoid growth and morphology over time.

### In vitro CRE recombinase transduction

For in vitro delivery of CRE recombinase, Takara’s CRE recombinase gesicles (Takara Bio USA, (Catalog No. 631449) were used according to the manufacturer’s instructions. Briefly, cells were seeded as organoids in mouse organoid media. The next day, Cre gesicles containing media was replaced with old media. CRE recombination efficiency was assessed by flow cytometry for mCherry (fused to the gesicles) 48 hours post-transduction.

### In vivo induction of neoantigens

Gp33 (neoantigen for CD8 T cells) and gp66 (neoantigen for CD4 T cells) were induced *in vivo* by administering 5 daily doses of doxycycline (50 mg/kg) and tamoxifen (80 mg/kg), as previously described (*17*).

### Histology and hematoxylin and eosin (H&E) staining

Tissue specimens were fixed in 10% neutral-buffered formalin, processed, and embedded in paraffin. Paraffin blocks were sectioned at a thickness of 4–5μm. Sections were stained with hematoxylin and eosin (H&E; Stat Lab, McKinney, TX) using standardized histological procedures.

### Triple immunofluorescence staining

Paraffin-embedded, 10% NBF-fixed mouse XG and prostate tissues were baked for 30 minutes at 70°C before loading onto the stainer. All staining was performed on a VMSI Discovery Ultra staining platform using the Ventana RUO DISCOVERY Universal procedure. For CD8a, CD4 and EpCAM triple immunofluorescence staining, **t**issues underwent two rounds of CC1 retrieval for 64 minutes each at 95°C, for antigen retrieval. An online DAPI counterstain was performed after the first CC1 incubation. After the second CC1 retrieval, a Disc Inhibitor was applied, followed by incubation with CD8a primary antibody, at 2.5μg/mL in casein diluent for 60 minutes at 37°C. A Goat Block was applied for 4 minutes, followed by Omnimap anti-Rabbit HRP for 16 minutes, and Rhod6G for 12 minutes. Slides were then subjected to denaturation with CC2 at 100°C for 8 minutes, followed by a neutralization step. Next, CD4 primary antibody was applied at 2.5μg/mL in casein diluent for 60 minutes at 37°C. This was followed by Omnimap anti-Rabbit HRP for 12 minutes and Cy5 for 8 minutes. A CC2 retrieval was performed for 8 minutes at 100°C, followed by a neutralization step. Next, EpCAM primary antibody was incubated at 0.123μg/mL in casein diluent for 32 minutes at 37°C. A Goat Block was applied for 4 minutes, followed by Omnimap anti-Rabbit HRP for 4 minutes, TSA-DIG for 8 minutes, and OP780 for 32 minutes. An offline DAPI counterstain was performed to improve nuclei counterstain for image analysis. Finally, sections were manually coverslipped with Prolong Gold Antifade mountant.

#### Fluorescent dyes

DISCOVERY Rhodamine 6G Kit (Ventana, Catalog #: 760-244), DISCOVERY Cy5 Kit (Ventana, Catalog #: 760-238), Opal 780 Reagent pack (Akoya, Catalog #: FP1501001KT). Opal TSA-DIG Reagent was reconstituted in DMSO (stock solution) and diluted 1:100 in 1X Plus Amplification Diluent (Akoya, Catalog #: FP1609) for a working solution. Opal 780 Reagent was reconstituted in ddH2O (stock solution) and diluted 1:25 in Blocking/Ab Diluent (Akoya, Catalog #: ARD1001EA) for a working solution.

#### Other Reagents

DISCOVERY Inhibitor (Ventana, Catalog #: 760-4840), OmniMap anti-Rb HRP (Ventana, Catalog #: 760-4311), VENTANA Goat IgG Block (Ventana, Catalog #: 760-6008), DAPI (Invitrogen Inc, Catalog #: D1306), Prolong Gold Antifade mountant (Molecular Probes/Invitrogen, Catalog #: P36930), DISCOVERY Wash (Ventana, Catalog #: 950-510), ULTRA LCS (Ventana, Catalog #: 650-210), Reaction Buffer (Ventana, Catalog #: 950-300), DISCOVERY CC1 (Ventana, Catalog #: 950-500), Ultra CC2 (Ventana, Catalog #: 950-223).

### Immunohistochemistry (IHC)

For Ki-67 and androgen receptor (AR) IHC, tumor and tissue samples were fixed in 10% neutral buffered formalin and paraffin-embedded. IHC was performed on 4μm-thick paraffin-embedded sections. Sections were de-paraffinized using xylene and rehydrated through graded alcohols to deionized water. For Ki-67, sections were treated with Target pH 6.1 antigen retrieval buffer (Agilent, Cat. No. S169984-2), and for AR, sections were treated with Target pH 9 antigen retrieval buffer (Agilent, Cat. No. S236784-2). Endogenous peroxidase activity was quenched using 3% hydrogen peroxide for both. Subsequent steps were performed by the Thermo Scientific Lab Vision Autostainer 360 platform at room temperature. Ki-67 antibody (Clone D3B5, Cell Signaling Technology, Cat. No. 9129) was incubated at 0.334μg/mL for 60 minutes following protein blocking with 3% bovine serum albumin. AR antibody (Clone EPR1535(2), Abcam, Cat. No. ab133273) was incubated at 0.1μg/mL for 60 minutes following protein blocking with 10% normal donkey serum in 3% bovine serum albumin. Detection of Ki-67 was performed using PowerVision polymer anti-rabbit HRP (Leica Biosystems, Cat. No. PV6119) for 30 minutes, and detection of AR was performed using the VECTASTAIN Elite ABC-HRP Kit (Vector Laboratories, Cat. No. PK-6100) for 30 minutes following Biotin-Donkey Anti-Rabbit IgG (Jackson Immuno Research Laboratories, Cat.No. 711-065-152). Visualization for both was achieved with 3,3’-diaminobenzidine (Thermo Fisher Scientific, Cat. No. 34065). Final steps were performed outside the Autostainer 360 platform. Sections underwent counterstaining with Mayer’s hematoxylin (Rowley Biochemical, Cat. No. L-756-1A), followed by dehydration through graded alcohols, clearing in xylene, and cover-slipping with a xylene-based mounting medium (Sakura, Cat. No. 6419).

### Flow cytometry

Single-cell suspensions were prepared from orthotopic and subcutaneous tumors with mechanical dissociation and collagenase IV digestion, followed by filtration through a 70μm cell strainer to obtain a single-cell suspension. Cells were then washed twice with cold PBS containing 2% FBS (FACS buffer). For surface staining, cells were first incubated with Fc Block for 10 minutes to minimize non-specific antibody binding and then incubated with a cocktail of fluorochrome-conjugated antibodies diluted in the FACS buffer for 30 minutes at 4°C, in the dark. Antibodies used were: anti-mouse PE-CD8 (catalog 561950), APC-H-2Db/LCMV gp33 (C9M) (KAVYNFATM) MHC Tetramer-Class I (catalog MHC-LC527), anti-mouse FITC-PD1 (catalog 135214). Following staining, cells were washed twice with FACS buffer. Samples were acquired on a BD LSRFortessa X-20 flow cytometer. Flow cytometry data were analyzed using FlowJo™ software.

### Immunoblotting

Organoids were isolated from the Matrigel basement membrane extract by trypLE (trypsin) treatment and washed 3 times with ice cold PBS in a 15-ml Falcon tube. Cells were lysed on ice in RIPA buffer containing protease inhibitors (Calbiochem) and phosphatase inhibitors (Calbiochem) and sonicated three times for 30 s at 30-s intervals using a Bioruptor. Lysate were centrifuged for 10 min at 20,000G at 4deg. Supernatant was collected as protein lysate. Protein concentrations were quantified using a bicinchoninic acid (BCA) assay (Pierce, ThermoFisher). Lysates were denatured using 4X protein loading dye (SDS 200 nM Tris, 8% SDS, 0.4% bromophenol blue, 40% glycerol, 400 mM 2-mercaptoethanol, pH 6.8). 10-20μg of protein was loaded on NuPage 4-12% gradient bis-tris polyacrylamide gels (Invitrogen). After electrophoresis, protein was transferred to a PVDF membrane and blocked with either 5% milk or 5% BSA in TBS-T. Primary antibodies were incubated overnight. Membranes were washed using TBS-T and incubated with secondary antibodies for 1 h at room temperature. Proteins were visualized using ECL and ECL prime (Amersham, GE healthcare) and ImageQuant 800 (Amersham, GE healthcare). Antibody concentrations and the catalog numbers are below. Blots were analyzed using FIJI/ImageJ software.

### Antibodies

Immunofluorescence and Immunohistochemistry: CD8a-Genentech’s internal antibody (Clone: 21E3.TRANS, Genentech, South San Francisco, CA). CD4 (Abcam, Catalog #: ab183685-1001), EpCAM (Abcam, Catalog #: ab32392). Naive Rabbit IgG (Cell Signaling Technologies (cst), Catalog #: 3900S) was used as an isotype control for all primary antibodies at a working concentration of 2.5μg/mL.

Immunoblotting: PTEN (cst 9559), TP53 (cst 2524), RB (cst 9309) and cyclophilin B (cst 43603).

### Spatial transcriptomics (Xenium) assay

Formalin-fixed, paraffin-embedded (FFPE) tissue blocks were sectioned at a 5μm thickness and placed on Xenium slides (10x Genomics). Slides were processed according to the manufacturer’s protocol, starting with deparaffinization and decrosslinking of the tissue sections (10x Genomics, CG000580 Rev E). A high-plex probe panel, combining a pre-designed 5,000-gene Mouse Pan Tissue and Pathway panel with a 100-gene custom panel (**Supplementary Table 1**), was hybridized to the target genes. Following the manufacturer’s protocol (10x Genomics, CG000760 Rev C), hybridized probes were subjected to rolling circle amplification generating localized and amplified signals for each targeted transcript. The slides were imaged, and the signal was decoded on a Xenium Analyzer (10x Genomics) to resolve transcript identities and their spatial coordinates.

### Zman-seq assay

Mice were anesthetized with isoflurane and intravenously injected with 100μl of fluorescent antibody solution with 4μg antibody corresponding to 0.2 mg/kg body weight, as previously described (*18*). Time stamping was done by injecting each mouse with a CD45 antibody solution at 36, 24, 12 h before euthanization. Antibodies (clone 30-F11) were labeled with AF647 for 36 h (BD Biosciences, Cat# 567376), PE for 24 h (Biolegend, Cat# 103106) and BV711 for 12 h (BD Biosciences, Cat# 563709). For harvesting prostate tumor tissue, animals were placed under general anesthesia with isoflurane. Thoracotomy with midline incision was performed, and the heart was exposed. A catheter was inserted into the ventricle for perfusion and the atrium incised to facilitate exsanguination. 10 ml of perfusate (1X sterile saline buffer) was flushed through the heart and the circulatory system. Finally, prostate tissue was harvested for mechanical and enzymatic dissociation of the prostate, and labelled immune cells were isolated with flow sorting. Three mice were used for each timepoint. Two biological replicates were performed for each timepoint.

Immune cells were fixed using the Parse Biosciences Low Input Cell Fixation v3 Kit (Parse Biosciences, Catalog number ECLC3303) and immediately barcoded with an Evercode Whole Transcriptome v3 Kit (Parse Biosciences, Catalog number ECWT3300). Briefly, scRNA-seq libraries were prepared using the Evercode Whole Transcriptome (WT) v3 kit (Parse Biosciences, Catalog number ECWT3300), following the manufacturer’s instructions, using a split-pool combinatorial indexing workflow (*66*).

### Patient prostate tumor scRNA-Seq data analysis

Publicly available scRNA-seq datasets (author provided raw counts) (author-generated raw counts) of prostate cancer disease progression were collected from the literature (*23–25*). Cell profiles with <500 detected genes or >20% mitochondrial counts and genes detected in <10 cells were filtered, yielding 104,861 cell profiles across 17,129 genes from 26 patients. Potential doublets were removed using Scrublet (v0.2.3) with default settings. Data integration was performed using Harmony [v0.2.0] with datasets as the batch variable. Unsupervised clustering using the Leiden algorithm was used to identify broad-level cell type populations, which were annotated based on canonical marker expression. T-cell state scores were calculated using published gene expression signatures (*67*) comprising naive/memory (35 genes), effector (37 genes), and dysfunction/exhaustion (64 genes) programs. Signature scores were calculated for each cell using Scanpy’s score_genes function, and cells were classified based on their highest scoring program.

### Cross-species projection of CAF subtypes

To validate mouse CAF subtypes in human patient samples, reference-query projection of human fibroblasts (n=5,351 cells) onto a mouse CAF reference atlas (n=268,380 cells from Xenium spatial transcriptomics) was performed. Mouse-human orthologs were identified using Ensembl BioMart (release 115), retaining one-to-one ortholog pairs. Of the 5,106 mouse genes, 4,538 (89%) had unambiguous human orthologs present in the patient scRNA-seq data. The mouse reference embedding was pre-computed during CAF subtyping: expression profiles were normalized (target sum 10⁴), log-transformed, scaled (max value 10), and reduced via PCA (50 components). Human fibroblast profiles were normalized and scaled identically, then projected into the mouse PCA space by matrix multiplication with the mouse PCA loadings across all 4,538 orthologous genes. UMAP coordinates for human cells were computed via weighted interpolation of the 10 nearest mouse neighbors in PCA space, with weights inversely proportional to Euclidean distance. CAF subtype labels were transferred from mouse to human using k-nearest neighbors (k=30) with majority voting, and prediction confidence was calculated as the fraction of neighbors sharing the majority label. Marker gene conservation was validated by comparing expression of known CAF subtype markers (COL1A1, THBS1, CCL2, LRRC15, PI16, DCN) across predicted human subtypes.

### Identification of malignant epithelial cells by copy number variation analysis

To distinguish malignant from non-malignant epithelial cells, copy number variation (CNV) analysis was performed using infercnvpy (v0.4.x). Gene coordinates were annotated using GENCODE v43 (GRCh38), and analysis was restricted to genes on standard chromosomes (chr1-22, X, Y). A sliding window approach (window size: 250 genes) was used to infer relative CNV profiles, using non-malignant cell types (B cells, T cells, myeloid cells, fibroblasts, and endothelial cells) as diploid reference populations. For each cell, a CNV score represents the magnitude of chromosomal aberrations genome-wide using the cnv.tl.cnv_score() function. To establish a data-driven classification threshold, multiple statistical criteria were evaluated based on the CNV score distribution of reference cells, including mean ± 2-3 standard deviations and 95^th^-99^th^ percentiles. CNV score distributions were compared between epithelial and reference cells with a Mann-Whitney U test. The 95^th^ percentile of reference cell CNV scores was selected as the classification threshold, which conservatively captures cell profiles exceeding normal chromosomal variation while accounting for technical noise. Epithelial cells with CNV scores above this threshold were classified as malignant. This approach identified widespread chromosomal alterations characteristic of prostate cancer, while minimizing false-positive classifications from diploid epithelial cells.

### Spatial transcriptomics data analysis

Raw data were processed using the 10x Genomics standard pipeline (xenium-4.0.0.19) including cell segmentation and the generation of gene-count matrices. Segmented cell profiles expressing fewer than 20 genes or containing fewer than 30 total counts were excluded, as were genes detected in fewer than 5 segmented cell profiles. Next, raw counts were preserved, and data were normalized by total counts per cell and log-transformed. The top 2,000 highly variable genes (HVGs) were identified for each sample, and their union was used for dimensionality reduction via Principal Component Analysis (PCA). A *k* nearest-neighbor graph (*k*=15) was constructed in PCA space, followed by UMAP visualization and clustering using the Leiden algorithm. All analyses were performed using Scanpy (v1.11.5) (*68*) with default parameters unless otherwise specified. Main cell types were annotated based on the marker genes (**fig. S2D**). Then, immune cells and fibroblast profiles were isolated, and a second round of iterative clustering (HVG selection, PCA, UMAP, and Leiden clustering) was performed using the same workflow to resolve finer subpopulations.

To analyze expression changes over time, basal cell profiles from wild-type control, 6-week post transplantation and 12-week post-transplantation were extracted. Pseuodobulk profiles were generated by aggregating gene counts per sample and converting to transcript per million (TPM), and analyzed by linear regression against numeric timepoints. Progressively up-regulated genes were selected based on regression slope >0 and p-value < 0.05. In addition, pair-wise differential expression analyses were performed between wild-type vs. 6-week post-transplant and 6-week post-transplant vs. 12-week post-transplant with Scanpy function sc.tl.rank_genes_groups. Gene Set Enrichment Analysis was performed using the python package GESApy with the database “GO_Biological_Process_2021”.

Spatial-aware cell-cell interaction analysis was performed using the python package stlearn function stlearn.tl.cci.run, and significant interaction pairs were selected based on FDR <0.05.

### Zman-Seq data analysis

Raw sequencing reads were processed using ParseBio’s trailmaker software [v1.6.0] to generate cell-count matrices. Cell profiles with >20% mitochondrial content were excluded. Clustering was performed using Scanpy’s [1.11.5,] (*68*). Gene expression counts were normalized to 10,000 reads per cell, log-transformed, and the top 2,000 highly variable genes were identified. Dimensionality reduction was performed using PCA. The top 50 principal components were used to construct a k-nearest-neighbor graph (*k*=15), which was used to generate a UMAP embedding. Clusters were defined using the Leiden algorithm. Cell annotation followed a hierarchical strategy based on marker genes (**fig. S4D**), as follows: T-lineage (*Cd3e*+), B-lineage (*Pax5*+), macrophages (*Csf1r*+), monocytes (*Ccr2*+), dendritic cells (*Flt3*+), neutrophils (*S100a9*+), and mast cells (*Gata2*+). Finally, sub-clustering was performed to define fine-grained cell states within Τ cells and macrophages: dysfunctional T cells (*Pdcd1*+), memory T cells (*Ccr7*+), T_regs_ (*Foxp3*+), iNKT cells (*Zbtb16*+), and NK cells (*Ncr1*+) (all from T cells) and M1-like (*Fn1*+, *Spp1*+), M2-like (*Mrc1*+), and antigen-presenting (*H2-Eb1*+) subsets (all from macrophages). For T_regs_, raw Treg counts were aggregated into a pseudobulk profile per sample, converted to transcript per million (TPM), and analyzed by linear regression against numeric timepoints. Gene Set Enrichment Analysis was performed with the Python package GESApy with the database “GO_Biological_Process_2021”. For CD8^+^ T cells, cell profiles were scored with Scanpy [1.11.5.]’s function sc.tl.score using established signatures (*67*) to identify memory and dysfunctional states genes. CellPhone DB v4.0 was used to infer ligand–receptor-mediated cell–cell communication at the 24-hour timepoint. To remove any sorting contaminants, immune cell profiles were subset from the full dataset, and raw counts for selected cell profiles were extracted. Data were normalized to 10,000 counts per cell and log-transformed using Scanpy. The processed expression matrix and corresponding metadata (sample ID, timepoint, and cell type) were used as input to the CellPhoneDB statistical analysis pipeline (*69, 70*), and significant (p-value < 0.05, permutation test) ligand–receptor interactions were identified for downstream visualization.

### Mouse tumor treatment studies

8-10 weeks old male C57BL/6J mice were subcutaneously injected with 10^6^ cells in 50ul, 25ul media + 25ul Matrigel. Two weeks following organoid injection, neoantigen expression was induced by administering 5 daily doses of doxycycline (50 mg/kg) and tamoxifen (80 mg/kg). Approximately two weeks after neoantigen initiation (*i.e.*, approximately 4 weeks post-organoid injection), mice were randomized into experimental groups based on established tumor volume. Following randomization, mice were treated for a period of four weeks. Tumor volumes were measured twice per week using digital calipers. At the experimental endpoint, mice were humanely euthanized by CO_2_ asphyxiation followed by cervical dislocation. Tumors were excised, weighed, and immediately processed for downstream analyses.

Anti-PD-1.mIgG2a LALAPG mAb (GNE 9899) and isotype control antibody, anti-gp120 mIgG2a (internally made at Genentech, PUR1CV43896) were administered IV at 10mg/kg on day 0 and then IP 3x/week for 4 weeks (for example, IP on day 2, 4, 7, 9, 11, 14, 16, 18, 21, 23, 25, 28). Ruxolitinib (INCB018424; Selleck Chemicals, Cat. No. S1378) was administered orally daily at 30mg/kg. Combination treatment was done by administration of anti-PD1 and Ruxolitinib, amount and mode of administration were identical to single agent dosing.

### eDOT Contrast

eDOT Contrast was performed as previously described (*71, 72*). Briefly, this calculation is a Dunnett contrast of endpoint Difference Over Time (eDOT) estimates. Such contrast of eDOT represents the difference in average growth rates between the treatment and reference groups that are calculated from the fits on the natural log scale over a common time period. This matches the actual slope of a fit on the natural log scale for tumors undergoing log-linear growth and corresponds to the “rise over run” of the given time interval for tumors undergoing non–log-linear growth. Mathematically, average tumor growth rates stem from integrating the first derivative of the fits on the natural log scale over the common study period, resulting in units of natural log units per day. The more negative the contrast value, the greater the antitumor effect; the more positive the contrast value, the greater the pro-growth effect. The 95% confidence intervals are based on the fitted model and variability measures of the data. Contrasts for which the 95% confidence interval does not include 0 are statistically significant with P<0.05.

### eGaIT Contrast

eGaIT Contrast was performed as previously described (*71, 72*). Briefly, this calculation is a Dunnett contrast of endpoint Gain Integrated in Time (eGaIT) estimates. Such contrast of eGaIT represents the difference in growth rates between the treatment and reference groups which are derived from the area under the curve (AUC) of the fits on the natural log scale over a common time period. To obtain the growth rate from the fit AUC on this scale in this time range, the AUC is corrected for starting tumor burden and then subjected to slope equivalence “normalization.” Mathematically, this “normalization” is attained by dividing the estimated baseline-corrected AUC value by half of the square of the common study period, resulting in units of natural log units per day. When tumors exhibit log-linear growth (i.e., the fit is a line on the natural log scale), slope equivalence “normalization” of the AUC results in the calculation of the slope of the fit. In cases where tumors demonstrate non–log-linear growth (i.e., the fit is curved on the natural log scale), slope equivalence “normalization” results in the calculation of the constant log-linear growth rate required to yield the observed baseline-corrected AUC for the fit. The more negative the contrast value, the greater the antitumor effect; the more positive the contrast value, the greater the pro-growth effect. The 95% confidence intervals are based on the fitted model and variability measures of the data. Contrasts for which the 95% confidence interval does not include 0 are statistically significant with P<0.05.

### Spatial Topological Domain Identification and Quantification

A multi-step spatial analysis pipeline was used to define topological domains based on malignant cell density and proximity. A malignant core niche was identified by a dual-constraint filtering approach. Spatial single cell profiles were assigned to the core if they were within 20 μm (*57*) of a basal malignant epithelial cell and within a high-density cluster, defined as having ≥10 malignant cells within a 50 μm radius (*56*). The malignant interface zone was defined the region within a 200 μm Euclidean distance from the malignant core boundary (*57*). All other regions were defined as non-malignant. The single-cell niche assignments were then transformed into continuous geometric representations by employing Delaunay triangulation to generate a concave hull around cell clusters, effectively merging individual coordinates into connected component polygon regions. T-cell infiltration was quantified by calculating the absolute density (cells/mm^2^) within these vectorized polygons. Statistical analyses of T cell density were performed to evaluate specific differences between individual treatment arms. *Post-hoc* pairwise comparisons were subsequently performed using Tukey’s Honestly Significant Difference (HSD) test, strictly controlling the family-wise error rate at 0.05 to account for multiple comparisons.

### Treatment interaction analysis

To test for interaction effects between treatments on cell proportion, a linear regression model was fit: Proportion ∼ PD1 + JAK + PD1×JAK, where PD1 and JAK are binary treatment indicators. The interaction term tests deviation from additivity: a significant positive coefficient indicates synergy (combination effect > sum of individual effects), a significant negative coefficient indicates antagonism, and a non-significant coefficient indicates additive effects (p > 0.05).

## Acknowledgments

We thank L. Gaffney and A. Hupalowska for their help in figure making and thank all members of the Regev lab for helpful discussions and feedback.

## Author contributions

A.N., and A.R. conceived the study. L.G. and A.R. jointly supervised this work. A.N. and A.R. wrote the paper, with input from all co-authors. A.N., H.W., Z.L. and A.R. designed the study, experiments and analysis and interpreted data. A.N., J.L., F.P., Z.A., J.L., S.G., S.C., J.J., S.R., L.M. and D.M. performed experiments and data interpretation. H.W., Z.L., R.J. and L.P. performed computational analysis of single cell data. K.C., E.L., M.S., B.D. and N.J. advised on the project.

## Data and code availability

Processed count matrices and cell metadata for 10X Xenium and Zman-seq can be downloaded at UCSC Cell Browser https://cells-test.gi.ucsc.edu/?ds=spatiotemporal-prostate-cancer.

Codes for data analysis are available for download at https://github.com/ZiyuLu041/spatial_prostate_cancer

---

**Supplementary Figure 1.**
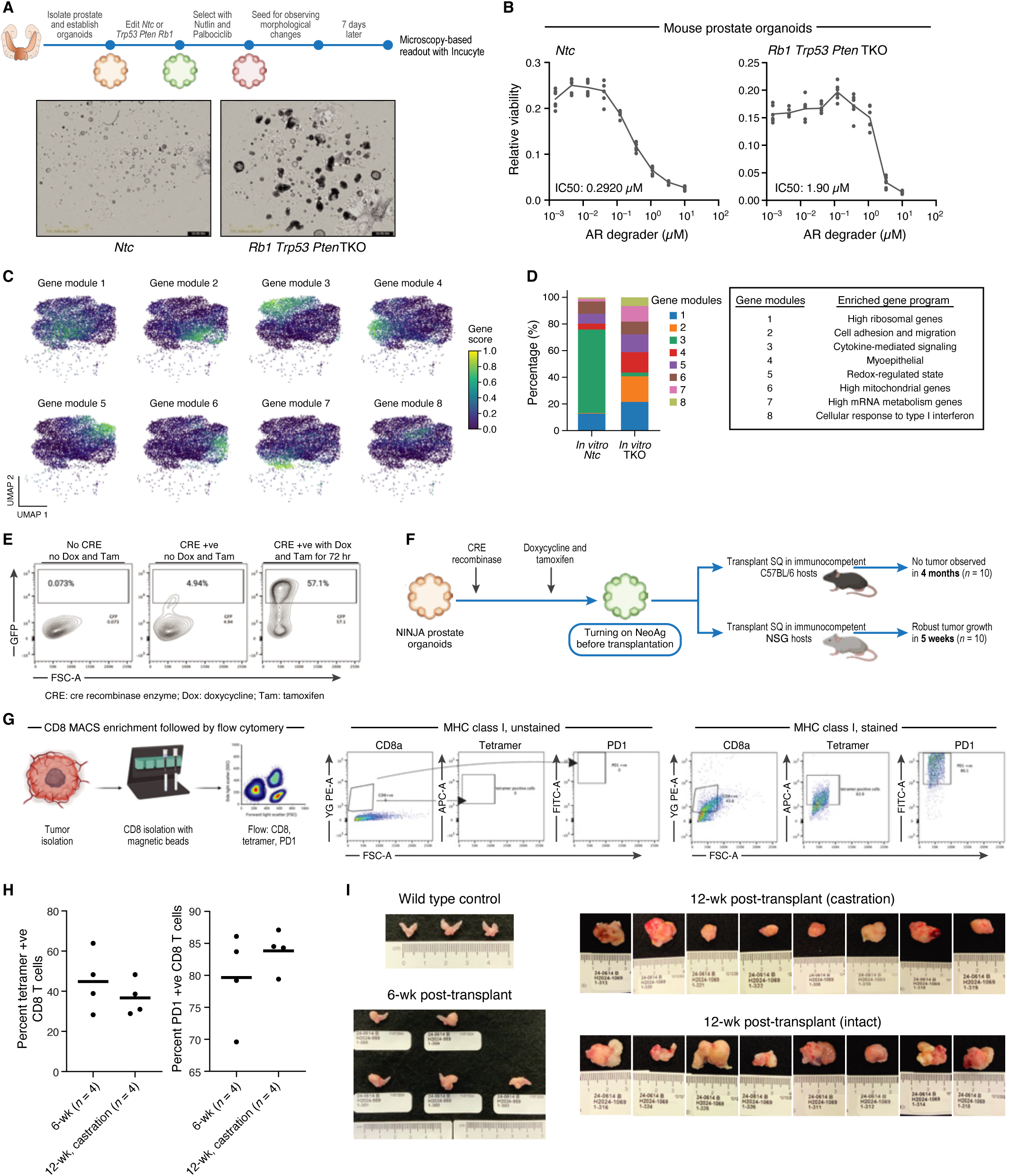
Characterization of genetically engineered prostate cancer organoids reveals differential therapeutic sensitivity and distinct cellular states. **A,** Experimental workflow for generating and assaying prostate organoid morphology. **B,** TKO organoids are more resistant to an AR degrader. Viability (y axis) of control (NTC, left) and TKO (right) organoids at different concentrations (x axis) of an AR degrader. IC50 values are shown at bottom left. **C,D,** Cell state heterogeneity in TKO organoids. C, UMAP embeddings of scRNA-seq profiles (dots) of NTC and TKO cells (as in Fig. 1A) colored by gene module scores. D, Fraction of cells (right) with top score for each gene module in control (NTC) and TKO organoids (x axis). Right: Enriched top GO terms (FDR<0.05 as in (*73*)) in each module. **E,** Neoantigen and GFP expression in NINJA prostate organoids. Flow cytometry plots of percentage of cells expressing GFP, in-frame with neoantigens, in NINJA prostate organoids without CRE recombinase (CRE), doxycycline, and tamoxifen (left), with CRE but without doxycycline and tamoxifen (middle), and with CRE, doxycycline and tamoxifen for 72 hours (right). **F**, Experimental mouse model. TKO organoids from NINJA mice treated with doxycycline and tamoxifen pre-transplantation were transplanted into immunocompetent (C57BL/6, n=10) and immunodeficient (NSG, n=10) mice. **G,H** CD8 T cell infiltration. G, (Left) Workflow. Right: Representative flow cytometry plots from stained (left) and unstained (right) samples, gated for CD8a, MHC class I tetramer (for neoantigen-specific T cells), and PD1 expression. H, Percentage (y axis) of neoantigen-specific (tetramer-positive, left) and PD1-positive CD8+ T cells (right), at 6- and 12-weeks post-transplantation with castration (x axis). Dots: individual mice. **I**, Prostates from different experimental groups.

**Supplementary Figure 2.**
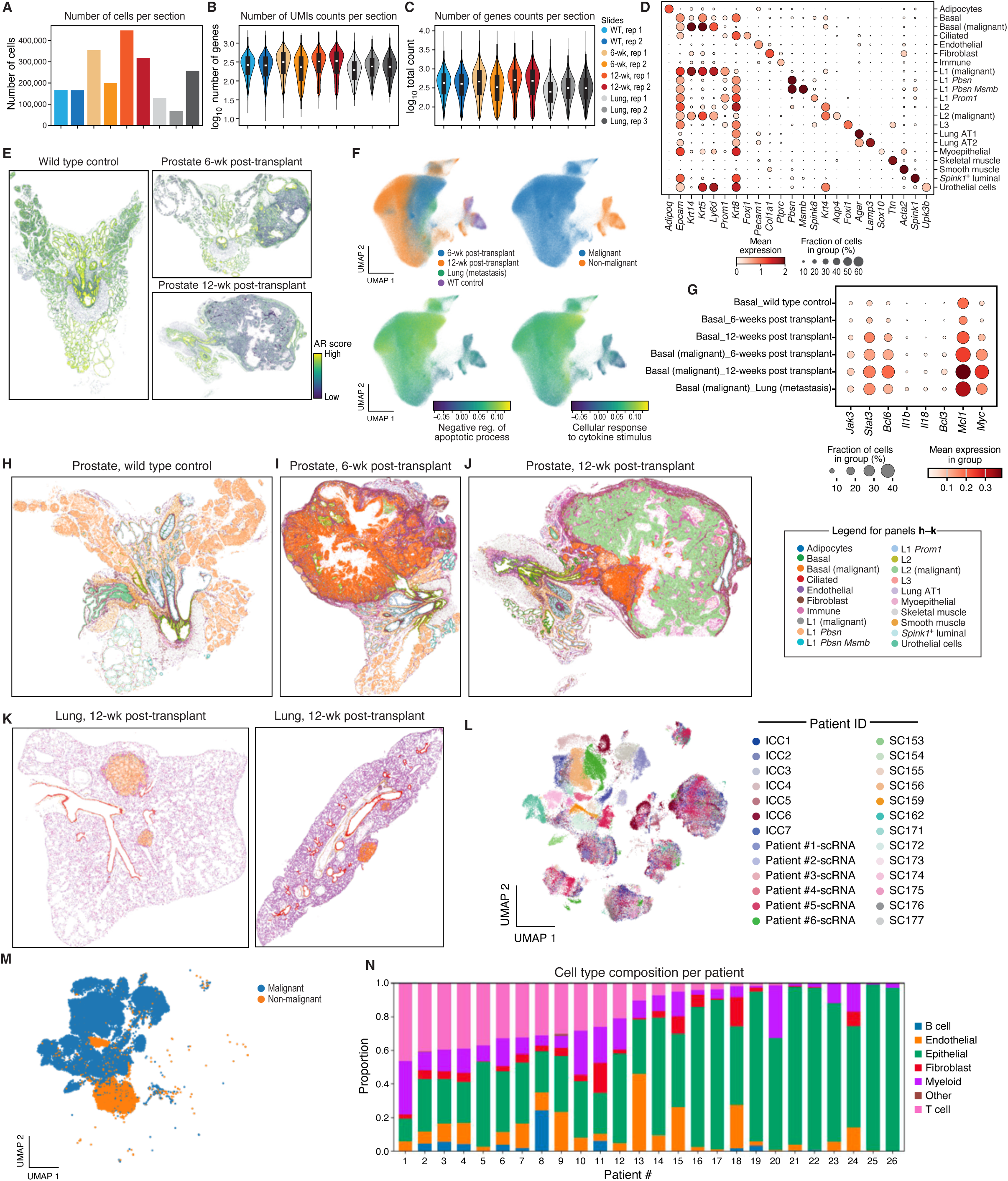
Spatial transcriptomics analysis of prostate cancer progression and metastasis. **A-C,** Quality controls. Number of cells per section (A, y axis), number (log10) of unique molecular identifier (UMI) counts per segmented cell (B, y axis), and number (log10) of genes detected per segmented cell (C, y axis) in each sample (x axis, legend). **D,** Marker genes for key subsets. Mean expression (dot color) and fraction of cells (dot size) expressing select marker genes (columns) in each cell subset (rows) based on expression profiles in segmented cells from spatial transcriptomics. **E,** Inferred AR activity in normal prostate and tumors. Spatial transcriptomics sections colored by the strength of an AR inferred activity score (color bar), calculated by expression of AR target genes in different prostate tissue (labels on top). **F,** Programs active in malignant prostate cells. UMAP embedding of segmented basal epithelial cell profiles (dots) colored by condition (top left), malignant status (top right), and select pathway scores (bottom). **G**, Marker genes for key basal epithelial cells across different conditions. Mean expression (dot color) and fraction of cells (dot size) expressing select marker genes (columns) in each cell subset (rows). **H-K,** Spatial transcriptomics maps of prostate and lung tissue. Spatial transcriptomics sections of control (top left), 6-wk (top middle) and 12-wk (top right) prostate tissue and of lung tissue with metastases (bottom) colored by segmented cell annotations. **L-N,** Integrated scRNA-seq analysis of human prostate tumors. L, UMAP embedding of scRNA-seq profiles (dots) from three prostate cancer studies (*23–25*) (as in Fig. 2K), colored by study-patient ID. M, UMAP embedding of prostate epithelial cells from the same studies (dots) colored by malignant state, as based on inferred copy number variation (CNV). N, Proportion (y axis) of cells in each major compartment (color legend) in each patient sample (x axis).

**Supplementary Figure 3.**
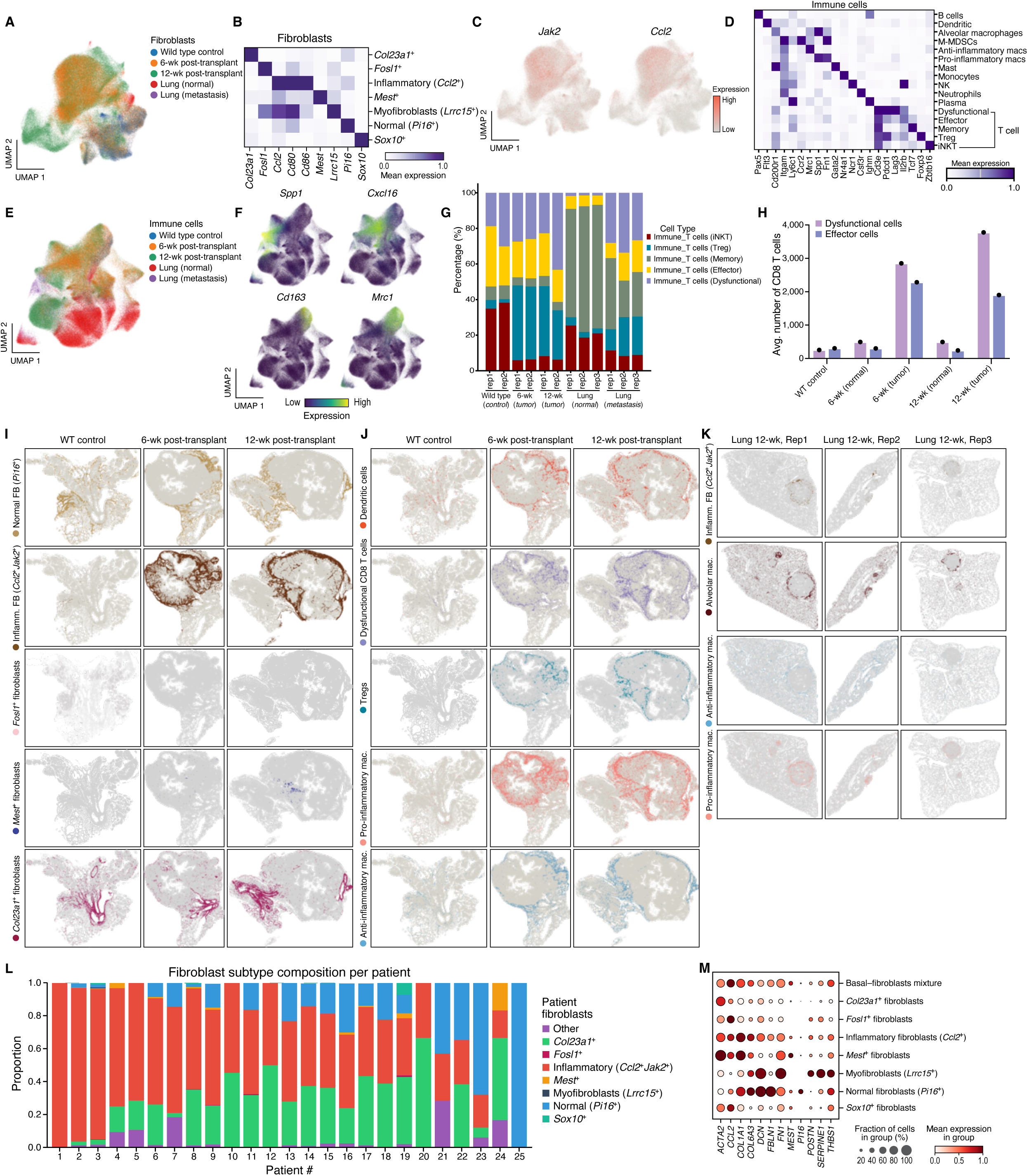
Spatial characterization of the TME. **A-C,** TME fibroblasts. A,C, UMAP embedding of RNA profiles of segmented fibroblasts colored by condition (A), or expression of key markers of inflammatory fibroblasts (C). B, Mean expression (color bar) of key marker genes (columns) in segmented fibroblast profiles from each subset (rows). **D-H,** Immune cells. E,F, UMAP embedding of RNA profiles of segmented immune cells colored by condition (E), or expression of markers of key macrophage subsets (F). D, Mean expression (color bar) of key marker genes (columns) in segmented immune cell profiles from each subset (rows).G, Proportion (y axis) of segmented cell profiles from each T cell subset in each tissue region, from each animal (x axis). H, Mean number of dysfunctional (purple) and effector (blue) CD8 T cells per region (y axis) in each condition (x axis). **I-K,** Spatial organization of key TME cell subsets. Spatial transcriptomics sections of prostate tissue from control (I, J, left) and 6wk (I, J, middle) and 12 wk (I, J, right) post-transplant or from lung metatases (I) transplantation colored by segmented cell annotations of different fibroblast (I), immune cell (J) or macrophages and fibroblasts (K) subsets. **L,M,** Fibroblast subsets in human prostate cancer. L, Proportion (y axis) of major fibroblast subsets identified in scRNA-seq (*23–25*) (*n*=25, data as in Fig. 2K) in each patient tumor (x axis). M, Mean expression (dot color) and proportion of expressing cells (dot size) of marker genes (columns) of different fibroblast subsets (rows) in scRNA-seq of human prostate cancer.

**Supplementary Figure 4.**
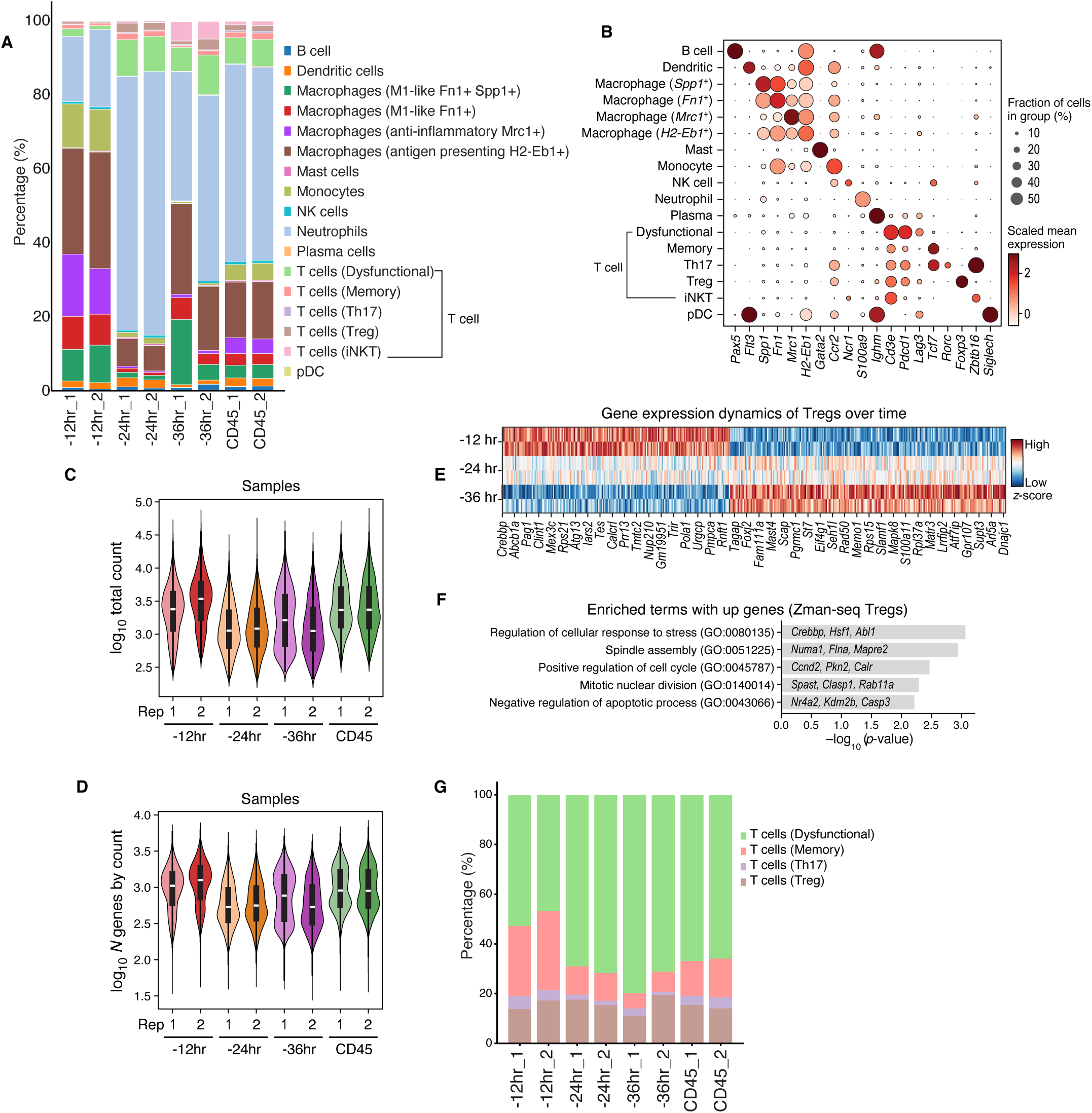
Quality control and cell type annotation of Zman-seq data. **A, B,** Cell type annotations. A, Percentage (y axis) of annotated immune cell subsets at different exposure times and overall in the tumor (x axis), in each biological replicate. B, Mean expression (dot color) and proportion of expressing cells (dot size) of marker genes (columns) of different cell subsets (rows) in Zman-Seq data. **C, D** Quality controls. Distribution of number of UMIs (c, log10, y axis) and number of genes (d, log10, y axis) per sample in each sample and fraction measured by Zman-Seq (x axis). **E, F,** Expression changes in infiltrating T_regs_. E, Normalized mean expression (z-score) in T_regs_ of genes (columns) that differentially expressed over exposure time to the TME (rows). F, Significance (x axis, -log_10_(P-value)) of top pathways (y axis) enriched for genes differentially expressed in T_regs_ (as in E). **G**, T cell compartment composition. Percentage (y axis) of annotated T cell subsets at different exposure times and overall in the tumor (x axis), in each biological replicate.

**Supplementary Figure 5.**
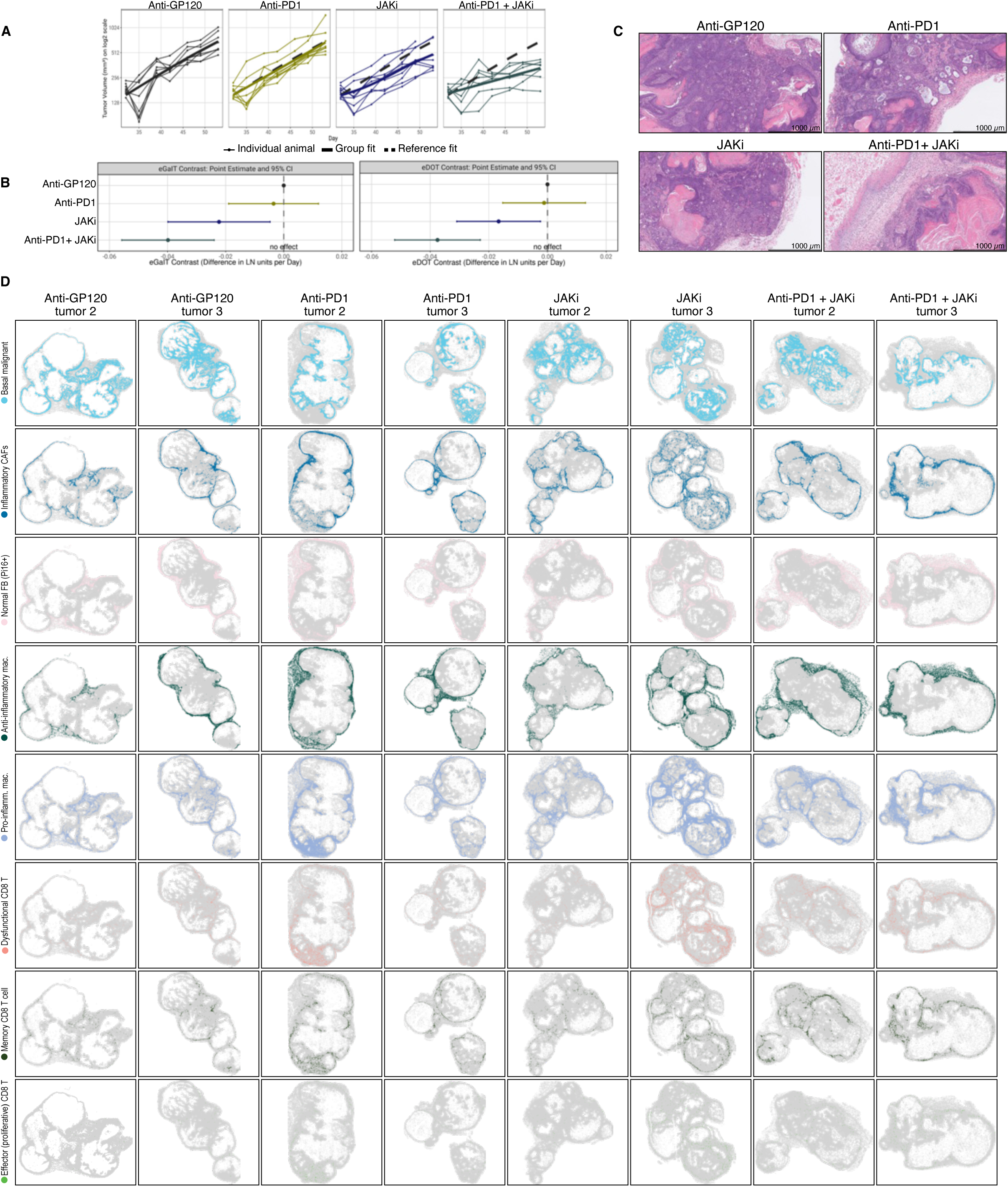
Changes in tumor growth and TME architecture in response to therapy. **A,B** Changes in tumor volume. A, Tumor volume (y axis, mm^3^) in individual mice (thin colored lines) over time (x axis) in each treatment group. Thick solid line: group fit; Thick dashed line: reference (anti-GP120) fit. B, Point estimates and 95% confidence intervals (CI) for treatment contrasts statistics (eGaIT (left) and eDOT (right), x axes) representing differences in tumor growth rates (natural log units per day). **C,D** Impact on histology and TME architecture. C, H&E stains of representative histology sections of subcutaneous TKO tumors from each treatment regime (label on top). D, spatial transcriptomics tumor sections from each condition (columns) colored by key cell subsets (rows).

**Supplementary Figure 6.**
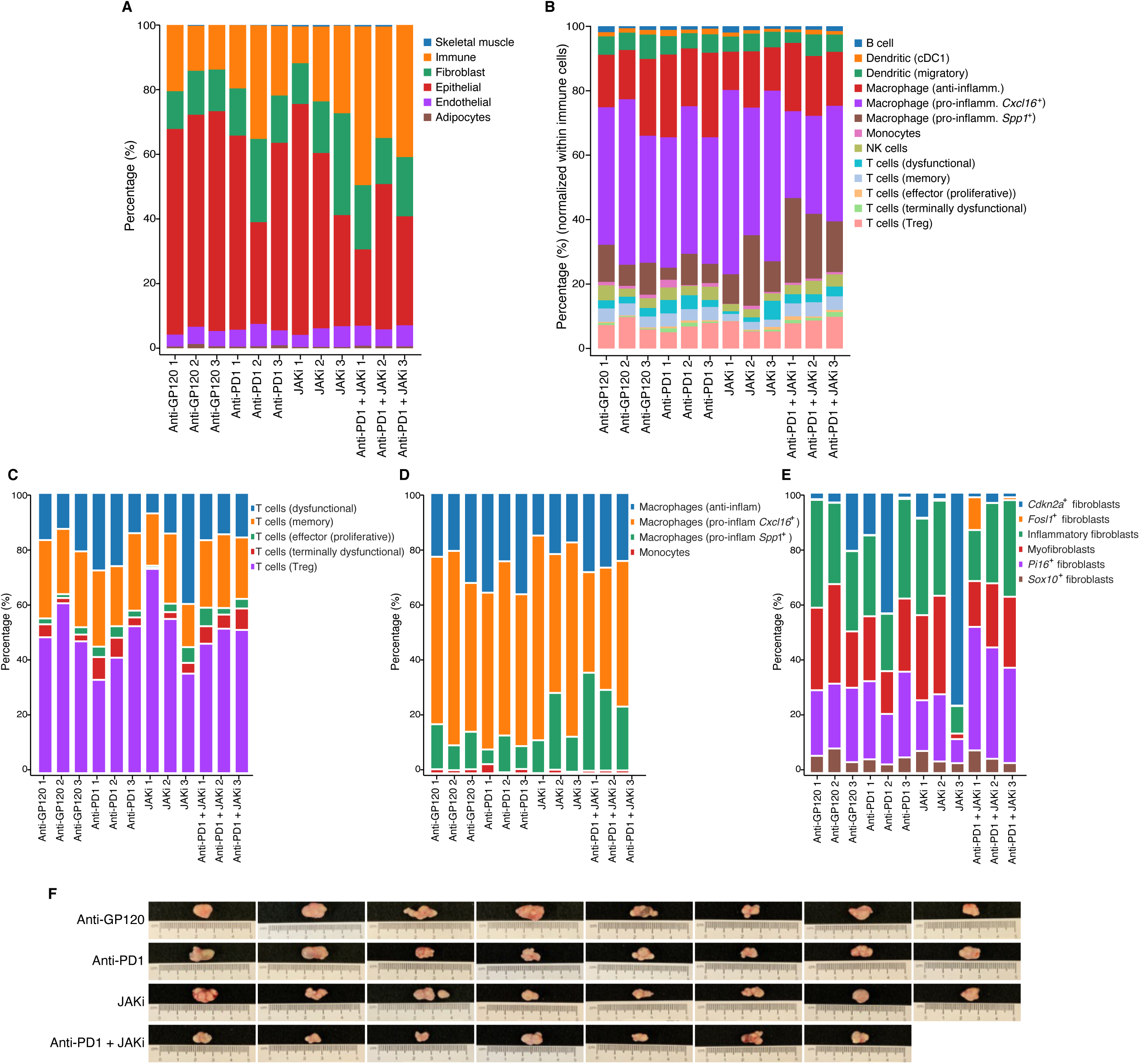
Cellular composition across individual tumors and treatment groups. **A-E,** Cell type proportion changes across treatment regimens. Percentage (y axis) of segmented cell profiles (color bar) in individual tumors from each treatment regimen (x axis). **F,** Macroscopic images of subcutaneous prostate tumors from each experiment in Fig. 5A.

**Supplementary Figure 7.**
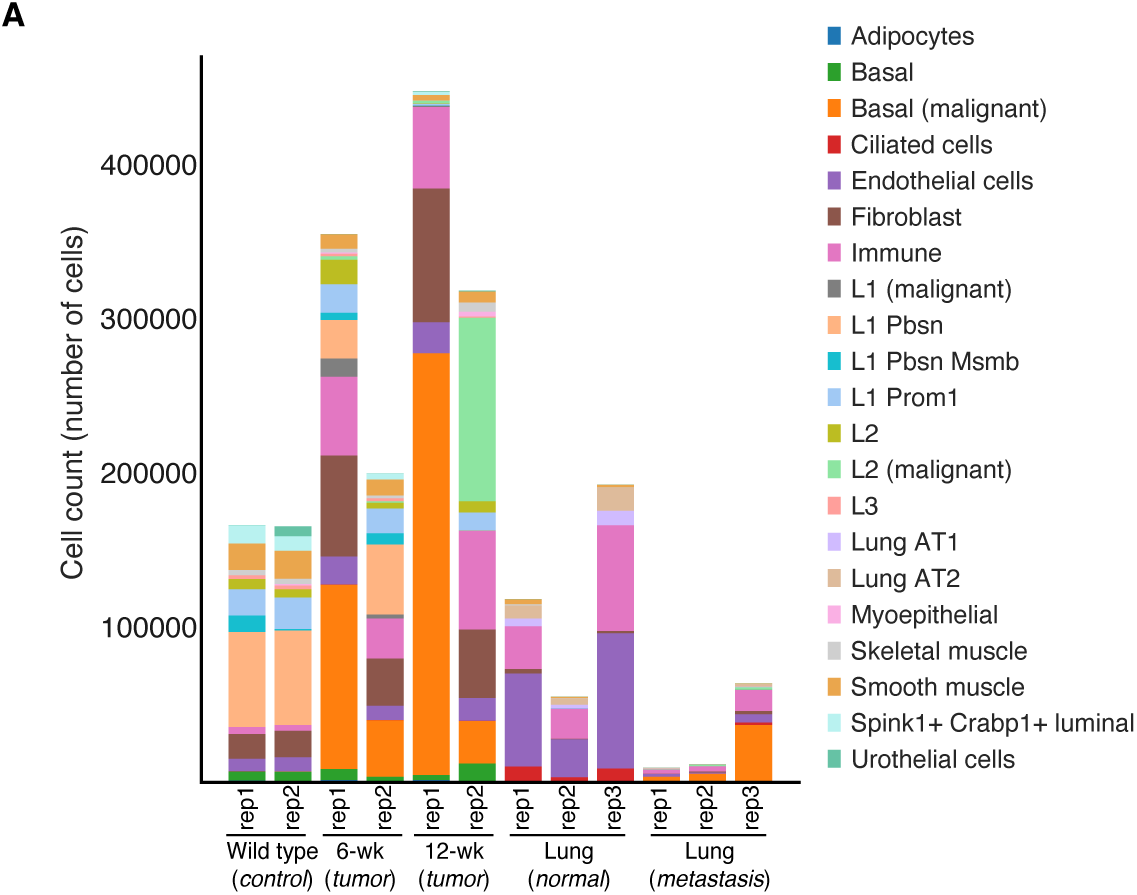
Composition of normal and tumor prostate and lung tissue. **A,** Cell number (y axis) in tumor and normal portions of each section in all animals from each time point (x axis), from experiment in **Fig 2B-D**.

